# Simple prerequisite of presequence for mitochondrial protein import in the unicellular red alga *Cyanidioschyzon merolae*

**DOI:** 10.1101/2024.02.19.581091

**Authors:** Riko Hirata, Yuko Mogi, Kohei Takahashi, Hisayoshi Nozaki, Tetsuya Higashiyama, Yamato Yoshida

## Abstract

Mitochondrial biogenesis relies on hundreds of proteins which are derived from genes encoded in the nucleus. According to characteristic properties of N-terminal targeting peptides (TP) and multi-step authentication by the protein translocase called the TOM complex, nascent polypeptides satisfying the requirements are imported into mitochondria. However, it has not been investigated whether the eukaryotic cell with a simple proteome and a single mitochondrion in a cell has a similar or simpler complexity of presequence requirements for mitochondrial protein import as other eukaryotes with multiple mitochondria. Based on the amino acid compositions of putative mitochondrial TP sequences in the unicellular red alga *Cyanidioschyzon merolae*, we designed the synthetic TP (synTP) and confirmed that synTP-fused mVenus were translocated into the mitochondrion *in vivo*. Through a series of experimental evaluations using modified synTPs, we showed that functional TP must have some basic residues, at least one, and compose the specific amino acid composition, but the physicochemical properties of net charge, hydrophobicity and hydrophobic moment are not strictly determined in *C. merolae*. Combined with the simple composition of the TOM complex in *C. merolae*, our results suggest that a regional positive charge in TP would be recognized and verified solely by TOM22 as a single-step authentication for mitochondrial protein import in *C. merolae*. The simple authentication mechanism indicates that the *C. merolae* cell, with its simple cell structure and genome, would not need to increase the cryptographic complexity of the lock-and-key for mitochondrial protein import.

## Introduction

Evolved from a free-living α-proteobacterial ancestor via an endosymbiotic event, the mitochondrion has their own genome and gene expression system (Gray et al., 1999). However, most genes encoding mitochondrial proteins have been now encoded in the nuclear genome DNA and 99% of mitochondrial proteins are synthesized by cytosolic ribosomes (Chacinska et al., 2009; Sickmann et al., 2003). Due to the compartmentalization by membranes, the nascent mitochondrial precursor proteins are secreted into mitochondria according to the embedded information in their peptide sequence (Pfanner et al., 2019; Schmidt et al., 2010). Except for mitochondrial membrane proteins, the signature for destination into mitochondria could be found in the N-terminal peptide sequence of precursor proteins called the presequence or targeting peptide (TP) (Endo and Yamano, 2009). The precursor proteins which have the claim are carried into the mitochondrion through the mitochondrial protein translocator of the outer membrane (called the TOM complex) (Araiso et al., 2019; Shiota et al., 2011; Shiota et al., 2015; Su et al., 2022). And then, precursor proteins are imported into matrix by the function of the TIM23 complex (Sim et al., 2023; Yamamoto et al., 2002; Zhou et al., 2023). The mitochondrial protein import system is widely conserved through the eukaryotes, suggesting it evolved in their last common ancestor.

To distinguish the nascent proteins which should be secreted into the mitochondrion from others, the mitochondrial targeting sequence has several characteristic features. The mitochondrial TPs contain usually 20 to 60 amino acids and tend to fold into amphiphilic α-helices (Calvo et al., 2017; Carrie et al., 2015; Gavel et al., 1988; Huang et al., 2009). The amino acid composition of them is generally enriched in alanine, leucine, lysine, and arginine. In particularly, it is considered that the presence of arginine residues and resultant a positive charge in the presequence determines mitochondrial targeting (Vögtle et al., 2009; von HEIJNE et al., 1989). The reason for the overall enrichment of basic amino acids is thought to be that the positive charge of TP facilitates passage through the electrochemical gradient across the inner mitochondrial membrane generated by the mitochondrial electron transport chain (Garg and Gould, 2016).

Despite these features are well found in the mitochondrial targeting sequences throughout eukaryotes, the common motif has not been identified and amino acid sequences differ from each other even in the same organism. Although the protein targeting sequence seems to be a cryptic sequence, significant biochemical features of the presequence indicate that destinations of each organelle protein are controlled and follow the uncharacterized rules in the eukaryotic cell. Given that each organism shows different trends of the mean length and amino acid composition of the presequences, there are countless functional sequences as the presequence and the complexity of the presequences might correlate with the complexity of cell structure such as multicellularity.

In this study, to understand the prerequisite of the TP for mitochondrial targeting, we experimentally assessed protein targeting function of various types of synthetic TPs using unicellular alga *Cyanidioschyzon merolae*. The *C. merolae* cell contains only one mitochondrion, which is easily identified by its shape and intracellular localization by fluorescence microscopy (Kuroiwa, 1998; Matsuzaki et al., 2004; Nozaki et al., 2007) (Fig. 1A). In addition, established gene targeting techniques using fluorescent reporters can be used to test whether the engineered TP has the ability to target a fluorescent protein (FP) or FP-fused protein to the mitochondria of *C. merolae* (Fujiwara et al., 2015; Imamura et al., 2009; Ohnuma et al., 2008; Tanaka et al., 2021). Through a series of *in vivo* and *in silico* experiments, we showed that an N-terminal peptide with the specific amino acid composition and very few basic residues fulfill the requirement for mitochondrial protein targeting. Thus, a key with a simple structure can open the mitochondrial protein gate in the monomitochondrial *C. merolae*.

**Figure 1.**
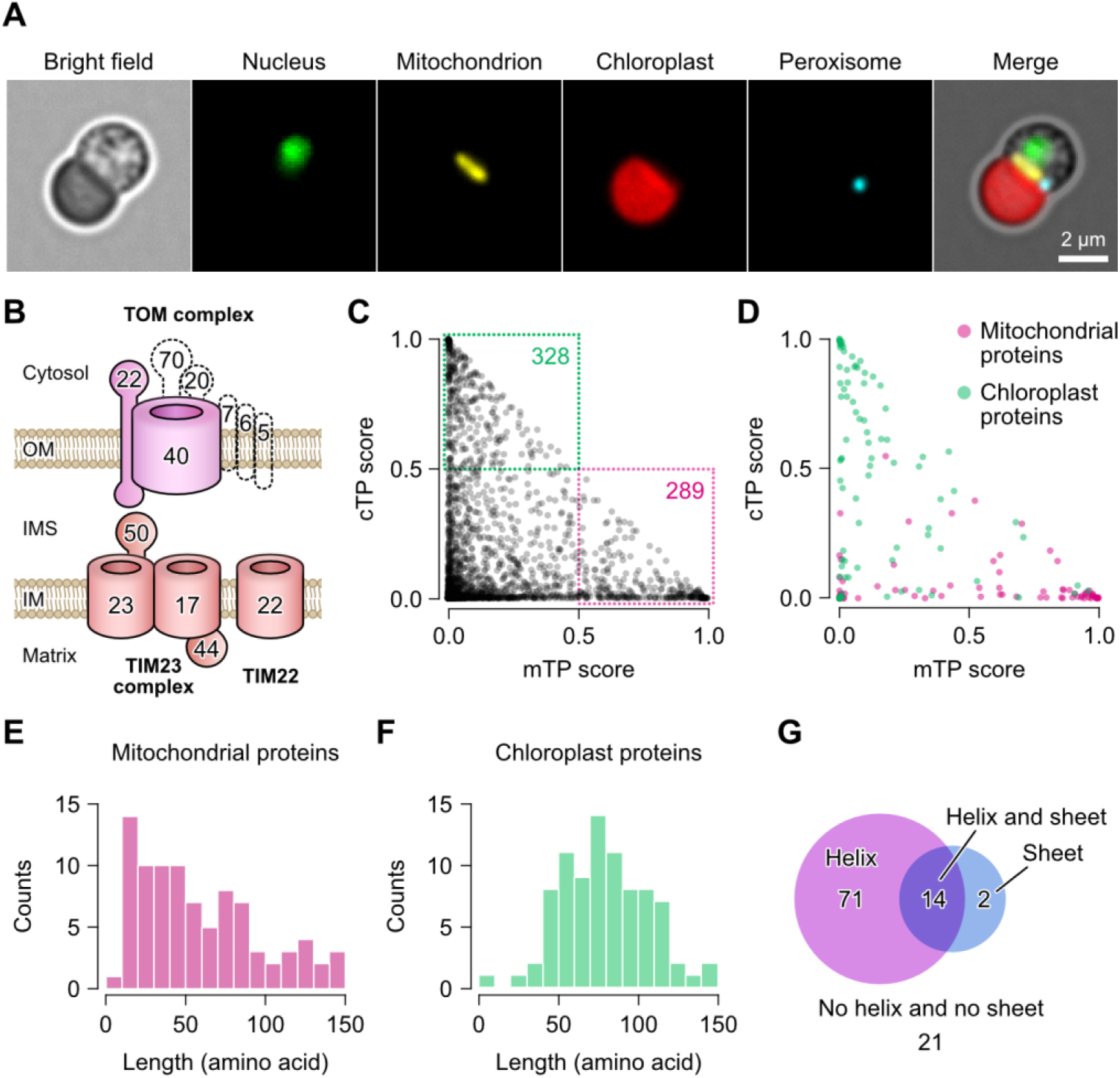
Genomic information of mitochondrial and chloroplast proteins in *C. merolae*. (**A**) Fluorescent images of a *C. merolae* cell. The *ACTIN* knockout cell was used for the imaging (Tanaka et al., 2021). Nucleus, mitochondrion, chloroplast, and peroxisome were visualized by Cas9-Venus, mScarlet, chlorophyll autofluorescence and mCerulean3. (**B**) The mitochondrial translocater of the outer mitochondrial membrane (TOM) complex and the inner mitochondrial membrane (TIM) complex. The TOM complex composes of β-barrel protein TOM40, α-helical membrane-integrated receptors TOM20, TOM22 and TOM70, and regulators TOM5, TOM6, and TOM7. Proteins which are not identified in the *C. merolae* protein-coding genes are illustrated with dashed lines. (**C** and **D**) Scatterplot comparisons of mitochondrial targeting and chloroplast targeting scores for all ORFs (4803 proteins) (**C**) and for well characterized mitochondrial (113 proteins) and chloroplast proteins (97 proteins) (**D**). Prediction scores for all ORFs are shown in Data S1 and the list of mitochondrial and chloroplast proteins is shown in Table S1 and S2. (**E** and **F**) Histograms of the length of presequences for the mitochondrion (**E**) and the chloroplast (**F**). (**G**) Venn diagram showing the classification of mitochondrial presequences containing α-helix (magenta), β-sheet (blue), both α-helix and β-sheet (purple), and neither α-helix nor β-sheet. See also Fig. S1 and S2.

## Results

### Genome analysis of the presequences in *C. merolae*

The fact that only 4803 genes are encoded in the nuclear genome and 99.9% of the protein-coding genes are single-exon genes suggests a simple proteome in *C. merolae* (Matsuzaki et al., 2004; Nozaki et al., 2007). Furthermore, as only two components, TOM40 and TOM22, in the TOM complex are characterized in the genome, the mitochondrial protein targeting system in *C. merolae* is likely to be functionally limited than that in other eukaryotic cells (Fig. 1B). To reveal primitive requisite as the functional TP for mitochondrial protein targeting, we first computationally evaluated all amino acid sequences encoded in nuclear genome by using a deep learning model-based presequence prediction tool, TargetP2.0 (Armenteros et al., 2019). By the analysis, 4803 open reading frames (ORFs) are classified into 336 of putative mitochondrial proteins (289 proteins over 0.5 mTP score), 349 of putative chloroplast proteins (328 proteins over 0.5 cTP score), and others (Fig. 1C; Data S1). To assess the predictability of the results, we confirmed the prediction scores for mitochondrial and chloroplast proteins that are well characterized in other organisms or experimentally confirmed to localize to the mitochondrion or the chloroplast in *C. merolae* (Mori et al., 2016; Moriyama et al., 2014a; Moriyama et al., 2014b) (Table S1 and S2). The recalls of protein targeting sequences for mitochondrial proteins (113 proteins) and chloroplast proteins (97 proteins) were 43.4% (49 proteins over mTP score 0.5) and 41.2% (40 proteins over cTP score 0.5), respectively (Fig. 1D). Given that the recall of presequence prediction for mitochondrial and chloroplast proteins in other organisms by TargetP2.0 was approximately 80-86% (Armenteros et al., 2019; Imai and Nakai, 2020), the lower prediction scores for *C. merolae* protein targeting sequences suggest that the mitochondrial targeting system in *C. merolae* not only shares basic similarities with those in other eukaryotes, but also has some differences.

To understand characteristics and principles of mitochondrial TP in *C. merolae*, we next compared the length of the putative mitochondrial and chloroplast TPs. In the comparison, TP regions in the mitochondrial and chloroplast proteins were presumed by protein sequence alignment analysis (see methods) and we omitted proteins whose length of putative TPs is shorter than 10 or longer than 150 amino acids from the analysis. The length of mitochondrial and chloroplast TPs are 59.3 ± 23.4 and 78.4 ± 17.3, respectively (Fig. 1E and 1F). Also, our dataset for mitochondrial TPs showed that 75.2% of mitochondrial TPs (85/113 proteins) are α-helical polypeptides (Fig. 1G; Fig. S1 and S2). Thus, as same with the results in other organisms, the mitochondrial TP is typically shorter than that of the chloroplast TP and the α-helical structure is the significant feature of the mitochondrial TP even in the *C. merolae* cell with the simplest organelle composition.

### *In vivo* analysis of mitochondrial targeting property by the fluorescence reporter assay

To verify whether the single α-helical polypeptide could work as a TP in *C. merolae*, an α-helix region (1-33 amino acids) or the full length (1-468 amino acids) of aspartate aminotransferase (AAT, CMC148C) was fused with a yellow fluorescence protein mVenus (Nagai et al., 2002) and introduced into the cells as a representative example (Fig. 2A and 2B; Fig. S3). Resultant transformants expressing either in AAT_1-33_-mVenus or AAT_full_-mVenus emitted fluorescent signals of the mVenus reporter in the mitochondrion. The result suggests that the α-helical polypeptide is one of the minimal requisites as a functional presequence for mitochondrial protein targeting in *C. merolae*.

**Figure 2.**
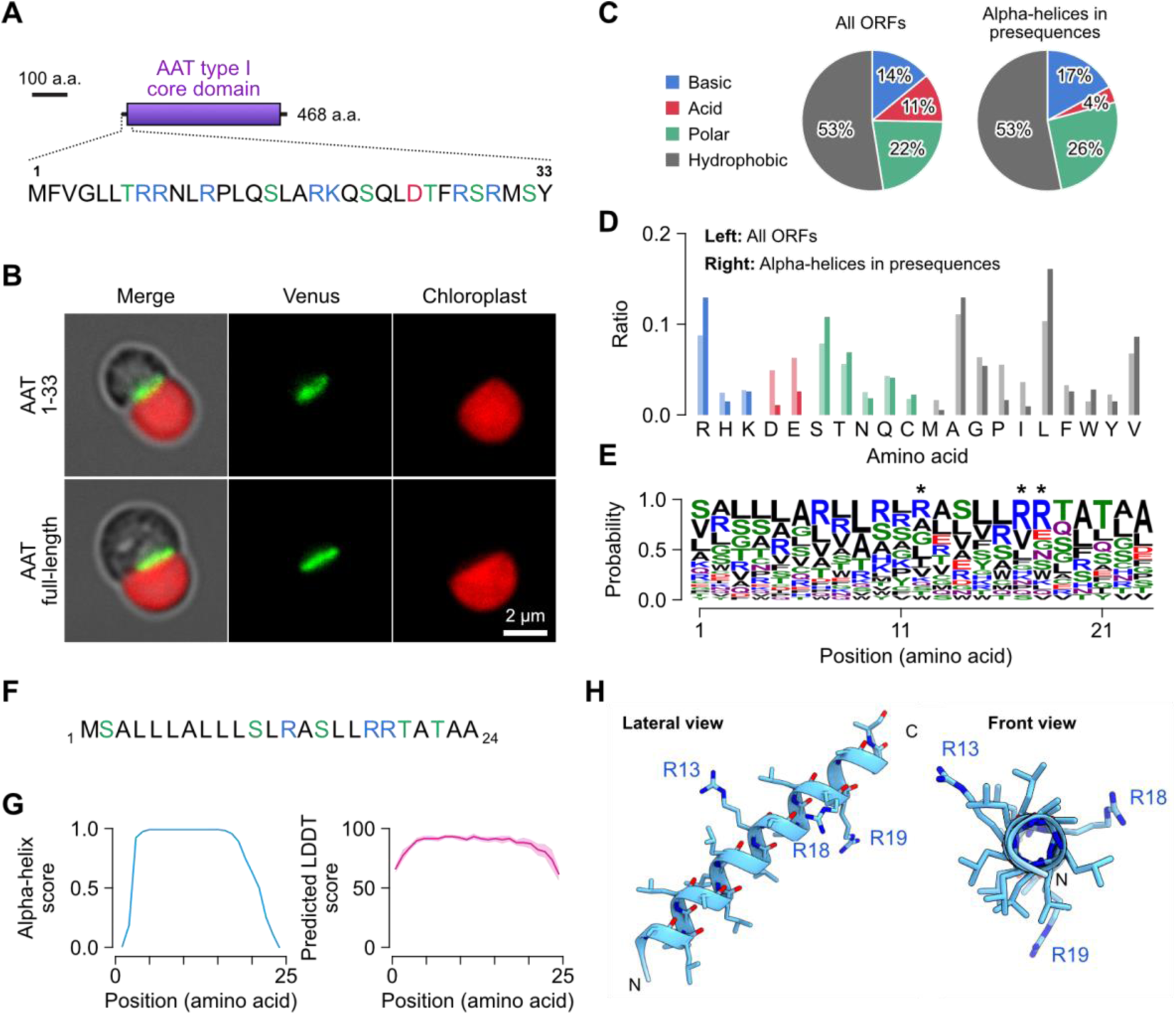
Synthetic mitochondrial presequence. (**A**) Scheme of 1-33 amino acid sequence of aspartate aminotransrferase (AAT, CMC148C). **(B**) Fluorescent images of AAT 1-33 and AAT full-length fused with mVenus. Fluorescent signals for mVenus are shown in green and chlorophyll autofluorescence are shown in red. See also Fig. S3. (**C**) Comparisons of amino acid compositions in all ORFs and α-helices in mitochondrial presequences. The α-helices in mitochondrial presequences are identified by structural simulation using the AlpahFold program. See also Table S3 for sequences. (**D**) Distribution of amino acids in all ORFs (left) and α-helices in presequences (right). (**E**) An amino acid sequence logo of α-helices in presequences. Asterisks indicate the arginine residues which are adopted in the synthetic presequence. (**F**) Sequence of the synthetic presequence of 24 amino acids. (**G**) Structural prediction score for α-helix (left) and local distance difference test score (right) of the synthetic presequence. (**H**) A simulated structure of the synthetic presequence in lateral and front views.

Next, we analyzed amino acid compositions of α-helices of the mitochondrial TPs. For the analysis, 31 single α-helical polypeptides, which are the length with less than 25 amino acids, in TPs has been evaluated (Table S3). As a result, we identified that the α-helical polypeptides compose of 17.3% of basic residues, 3.6% of acid residues, 26.2% of polar uncharged residues and 52.9% of nonpolar residues (Fig. 2C, right). Given that the average amino acid composition of all *C. merolae* ORFs is 11.2% of basic residues, 14.0% of acidic residues, 22.2% of polar uncharged residues and 52.5% of nonpolar residues (Fig. 2C, left), although the decrease in the proportion of acidic residues is remarkable, the overall trend is not extremely skewed in the mitochondrial TPs. More clear differences were found not in chemical characters, but in the amino acid compositions. While aspartic acid, glutamic acid, proline, and isoleucine residues are scarce, the compositions of arginine, serine, threonine, alanine, leucine, and valine residues are abundant in the mitochondrial TPs (Fig. 2D). In these abundant residues, it is known that alanine, arginine and leucine have high α-helical propensities (Pace and Scholtz, 1998). Since the mean hydrophobic moment in the α-helical polypeptides was 0.288 ± 0.13 μH (Table S3), the TP is very weak amphiphilicity. Also, due to the low composition of acidic residues, the average charge of the α-helices was estimated to be 2.80 ± 1.76 at pH 7.0 (Table S3). Thus, the mitochondrial TP has a specialized amino acid composition compared with other polypeptides and a tendency to form α-helical structure with weak amphiphilicity and weak positive charge in *C. merolae*.

### Mitochondrial targeting property of designed presequence

To further investigate the basal prerequisites of the mitochondrial TP, we designed a synthetic TP (synTP) by linking the most frequent amino acid residue at each position in the α-helices (Fig. 2E). In order to make the charge similar to that of the endogenous TPs, the two arginine residues have been replaced to leucine and serine residues, respectively. As a result, the synTP contains three arginine residues in the 24 amino acids (Fig. 2F). Computational simulations of both the secondary and tertiary structure indicated that the synTP forms a single α-helical structure (Fig. 2G and 2H). By introducing synTP-fused mVenus into the cell, we detected the fluorescent signal in the mitochondrion and concluded that the designed TP has the mitochondrial targeting property (Fig. 3, left). As it is well known that multiple basic residues in the TP are required for protein targeting into the mitochondrion (Gavel et al., 1988), we investigated the requirement of the minimum number of basic residues for mitochondrial protein targeting in *C. merolae*. To investigate this, arginine residues in the synTP were progressively replaced by other residues. Interestingly, while computational simulations predicted that modified synTPs containing less than 2 arginine residues would lose the ability to translocate to the mitochondrion (Table S4), transformants expressing modified synTP fused to mVenus showed that even a single arginine residue in the TP fulfills the role for mitochondrial targeting (Fig. 3). In addition, we confirmed that lysine residues are exchangeable for arginine residues in the TP (Fig. 3, right).

**Figure 3.**
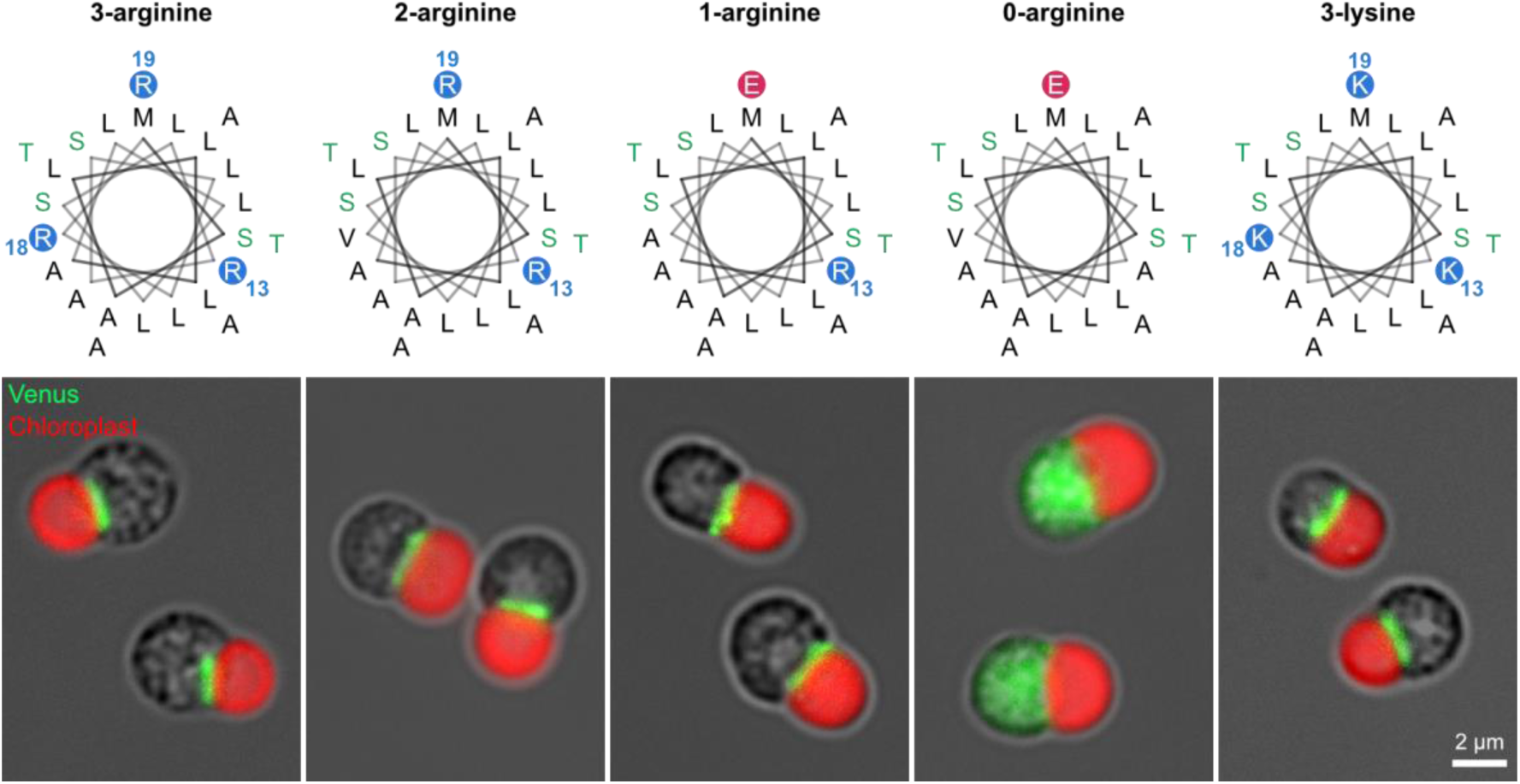
Effects of the number of basic residues in the synthetic presequence. The helical wheel diagrams for amino acid sequences of the synthetic presequences containing 3, 2, 1 or 0 arginine residues and 3 lysine residues. Representative images of cells are shown in the bottom of each helical wheel.

### Identification of physicochemical properties for functional presequence

Generally, mitochondrial TPs contain an arginine residue near the protease cleavage site with a sequence motif of R-X-/-X or R-X-F/T/L-/-A/S-X (Heidorn-Czarna et al., 2022), indicating that not only the net charge but also position of arginine residues would affect the mitochondrial targeting property. Based on the assume, we investigated the effect of the localization of the single arginine residue in the synTP^1R^ by arginine scanning approach. The position of a single arginine residue in +2, +6, +8, +9, +12, +13, +15 or +21 was examined (Fig. 4A). After the series of analyses, we found that the position of the single arginine residue in the synTP^1R^ is permissible broadly in the α-helix, but one of synTP^1R^ containing arginine residue at +2 position relative to the initial methionine residue lost the property and the mVenus reporter localized in the cytosol like synTP^0R^ (Fig. 4B). Taken together, an arginine residue in α-helix can give the functional property for the mitochondrial TP and the position of arginine residue is allowed in broad range of the helical structure except for the flanking position. By changing either the number of arginine residues or the position of the arginine residue, the physicochemical properties of synTP are drastically altered (Fig. 4C to 4E). Since the hydrophobicity and hydrophobic moment are acceptable in the broad range (0.388 to 0.747 H and 0.246 to 0.101 μH) as functional TPs, the importance of these factors for protein targeting to the mitochondrion is not high. Furthermore, neither the hydrophobicity nor the hydrophobic moment of the mitochondrial TP in *C. merolae* is characterized. More importantly, the *C. merolae* mitochondrial TP functioned normally even when the charge was negative. While the results suggest that the net charge of the TP does not need to be positive for protein targeting to the mitochondrion, a modified TP without an arginine residue (synTP^0R^) lost the ability to target mitochondrial proteins (Fig. 3). It is therefore suggested that basic residues in the α-helix, with the exception of the flanking position, are essential for a functional TP, but their importance is not linked to the net charge of the TP.

**Figure 4.**
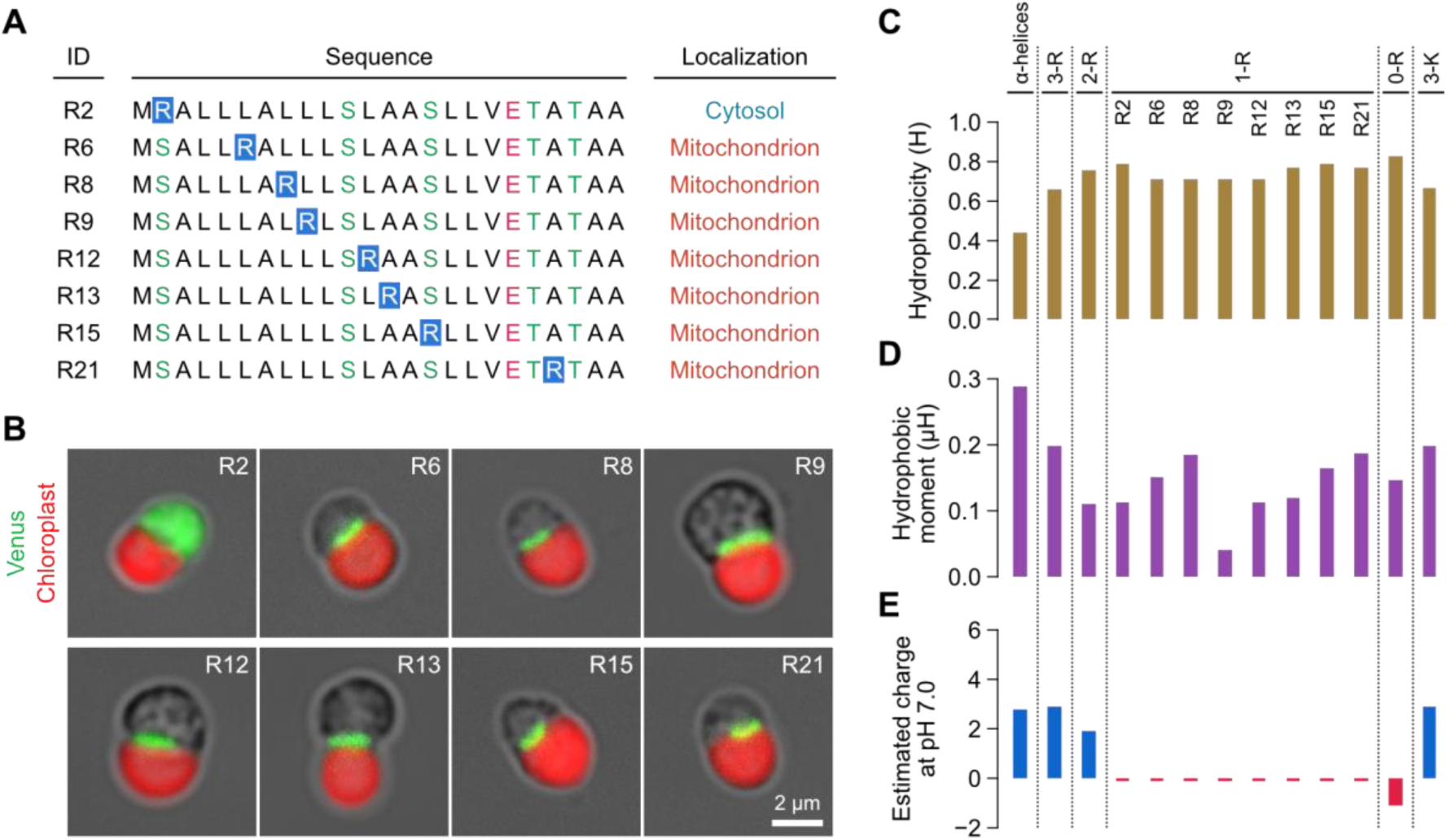
Arginine scanning analysis and physicochemical properties of the modified synthetic presequences. (**A**) Arginine scanning of the modified synthetic presequence. (**B**) Fluorescent images of each modified synTP^1R^-mVenus. (**C** to **E**) Physicochemical properties of each synthetic presequence. See also Table S4.

### Verification of mitochondrial targeting property of endogenous polypeptides sharing sequence similarity with the synTP

Our results showed that the specialized amino acid composition and the presence of a few basic residues in the α-helix are prerequisites for the functional TP in *C. merolae*, however it is questionable whether these lax requirements are sufficient to distinguish mitochondrial protein precursors from other proteins *in vivo*. If the peptide sequence of the synTP is very specific in protein sequences and only endogenous mitochondrial TPs have sequence similarity with the synTP in *C. merolae*, the sequence would act as a key for the mitochondrial protein gate. On the other hand, if the sequence has some similarity to other proteins, gene products with translation errors or genetic mutations that cause amino acid substitutions in each gene would easily lead to misdelivery of protein to the mitochondrion. To address this issue, we evaluated the sequence similarity between the N-terminal peptides (1-24 amino acids) of all ORFs and the synTP using the BLOSUM30 matrix.

Consequently, not only mitochondrial proteins but also many cytosolic and chloroplast proteins are highly scored (Table S5). For example, a hypothetical protein (CMD051C), a putative formin-like protein (FMNL, CMN049C) and a putative chloroplast ribosomal protein S1 (RPSA, CMM019C) were ranked first to third. The N-terminal peptides for FMNL and PRSA are also highly predicted to form an α-helical structure by local structure prediction, as is the case for synTP (Figure 5A and 5B). FMNL and chloroplast RPSA are well studied as a cytosolic protein related to actin filament dynamics and a 30S ribosomal protein S1 in chloroplasts, respectively, but it has not been reported that these proteins localize to mitochondria.

**Figure 5.**
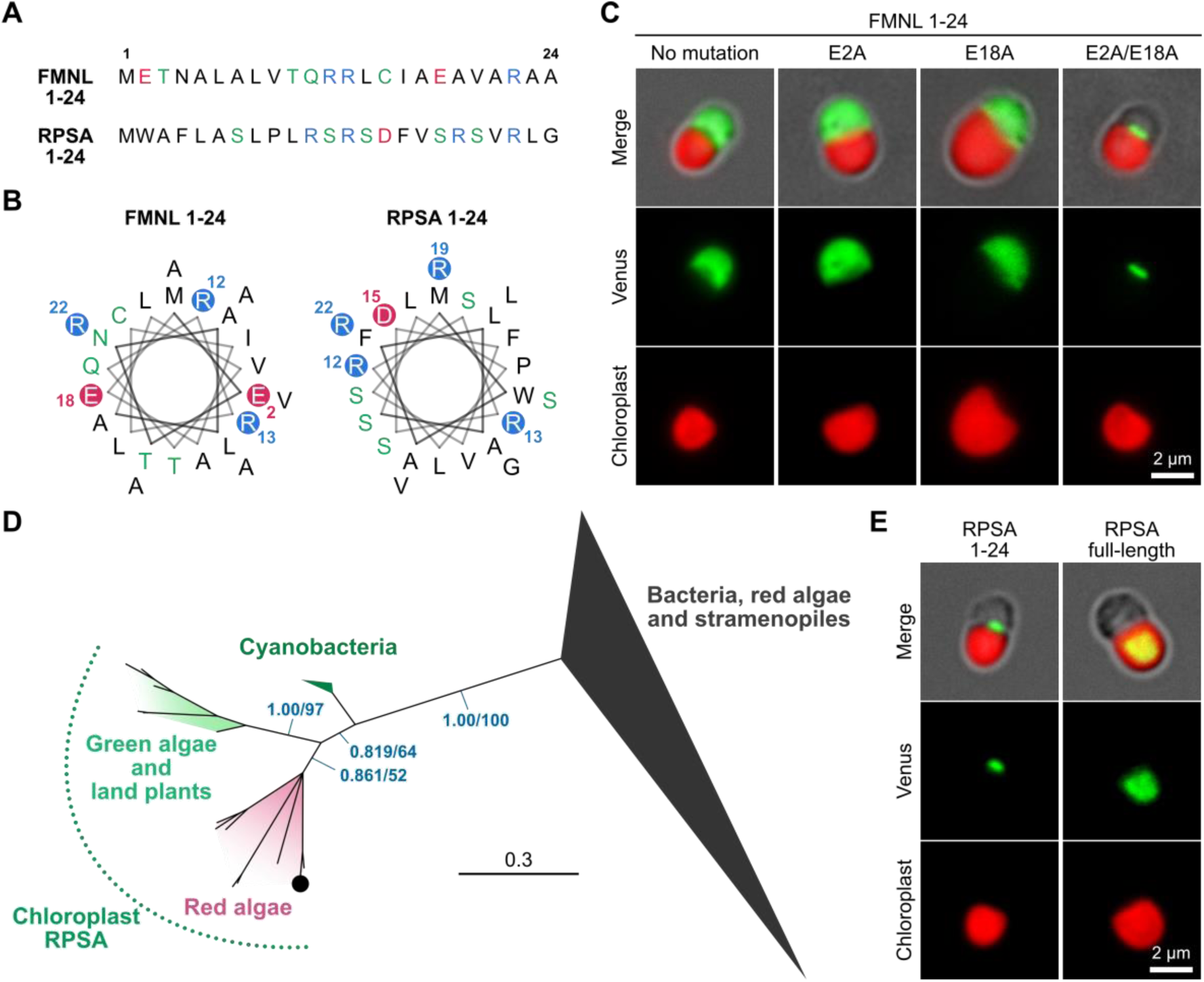
Fluorescent reporter assay for putative presequences of formin-like protein and chloroplast r-protein S1. (**A**) Sequences of formin-like protein (FMNL) and chloroplast r-protein S1 (RPSA) of 24 amino acids. (**B**) Helical wheels for FMNL 1-24 and RPSA 1-24. (**C**) Fluorescent images of the N-terminal 1-24 peptide of FMNL fused with mVenus. Mutated FMNLs (E2A, E18A and E2A/E18A) fused with mVenus are also shown. (**D**) The Bayesian tree of chloroplast, mitochondrial and bacterial RPSA proteins (See Fig. S4 for the unabbreviated tree). Numbers at left and right above branches indicate posterior probabilities of Bayesian inference and bootstrap values of the maximum likelihood method, respectively. Branche lengths are proportional to the evolutionally distances indicate by the scale bar. The *C. merolae* chloroplast RPSA (CMM019C) is shown as the black circle and the mitochondrial RPSA (CMI304C) is involved in the triangle. (**E**) Fluorescent images of the RPSA 1-24 or RPSA full-length fused with mVenus.

### Fluorescence reporter assay for the native or modified N-terminal polypeptides of FMNL

To determine whether these N-terminal peptides have mitochondrial targeting function, we performed the fluorescent reporter assay for the 1-24 amino acids of FMNL. As a result, the fluorescent signal for FMNL 1-24 fused mVenus was identified in the cytosol, suggesting that FMNL 1-24 is similar to synTP but does not have a mitochondrial targeting property (Fig. 5C, left). As a notable difference between FMNL 1-24 and synTP is the total number of acidic residues, we hypothesized that a negative charge derived from these residues prevents translocation of FMNL to the mitochondrion. Therefore, two acidic acid residues (E2 and E18) were progressively replaced by alanine residues. Using this approach, we found that a single substitution of glutamic acid residues did not affect the targeting property, but a double mutation (E2A and E18A) conferred the targeting property to the mitochondrion (Fig. 5C). This demonstrates that a small number of amino acid substitutions can alter destination of a protein *in vivo*.

### Evolutional origin and the targeting property of the N-terminus of RPSA

As a second example, we investigated the targeting property of the N-terminus of RPSA. Prior to the *in vivo* assay, we performed the phylogenetic analysis to verify whether the *C. merolae* RPSA CMM019C shares sequence similarity and is evolutionarily related to other chloroplast and cyanobacterial RPSAs. The results showed that cyanobacterial, green algal/land plant, and red algal RPSAs containing the *C. merolae* RPSA formed a phylogenetic group and were separated from other bacterial and eukaryotic RPSAs (Fig. 5D, Fig. S4). The evolutionary relationship between *C. merolae* RPSA (CMM019C) and cyanobacterial RPSA suggests that the gene product of CMM019C would function as 30S r-protein S1 in the chloroplast, not in the mitochondrion. Next, we tested the targeting property of the N-terminal peptide sequence of the chloroplast RPSA using the fluorescent reporter assay. As a result, we found that the N-terminal peptide has the property to translocate to the mitochondrion but not to the chloroplast (Fig. 5E, left). We also confirmed that the full-length RPSA fused to mVenus translocated to the chloroplast (Fig. 5E, right). The results suggest that the N-terminus of RPSA has the same targeting property to the mitochondrion as the synTP, but an additional peptide sequence would convert it to the chloroplast TP.

## Discussion

### The single-step authentication of mitochondrial preproteins in *C. merolae*

Through a series of *in vivo* and *in silico* experimental evaluations using synTPs, we showed that the minimal prerequisite for protein targeting to the mitochondrion in *C. merolae*. The results indicated that functional TPs need to have some basic residues, at least one, in an α-helix comprising the specific amino acid composition, but the physicochemical properties of the net charge, hydrophobicity and hydrophobic moment do not seem to be strictly determined in *C. merolae*. The reason for the small number of basic residues in the TP could be explained by the characteristics of the TOM complex in *C. merolae*. Previous pioneering studies in fungi and animals indicated that the mitochondrial outer membrane proteins TOM20 and TOM22 identify mitochondrial TPs by multi-step authentication and introduce mitochondrial precursors into the mitochondrial intermembrane space (Abe et al., 2000; Araiso et al., 2019; Hanif Sayyed and Mahalakshmi, 2022; Nargang et al., 1998; Pfanner et al., 2019; Saitoh et al., 2007; Shiota et al., 2011). During the process, precursor proteins in the cytosol are initially detected and captured by TOM20 on the mitochondrial outer membrane via the hydrophobic interaction between the hydrophobic groove on the TOM20 surface and the hydrophobic site of the amphipathic helix of TP (Abe et al., 2000; Saitoh et al., 2007). Then, TOM22 interacts with the TP by the electrostatic interaction between the negatively charged site of TOM22 and the positively charged site of the helix in the TP (Nargang et al., 1998; Shiota et al., 2011; Su et al., 2022). After the verification by TOM20 and TOM22, precursors are delivered to the β-barrel protein transporter TOM40 and the TIM complex. While a TOM22 homologue has been identified in *C. merolae*, there is no sequence encoding TOM20 in the genome (Fig. 1B). Given that hydrophobicity is not strictly required for the functional TP (Fig. 4C), combined with the absence of TOM20 in the genome, verification of the mitochondrial precursors in *C. merolae* would be performed by TOM22 alone as a single-step authentication.

Interestingly, the N-terminal domains of Human TOM22 (142 aa) and *C. merolae* TOM22 (119 aa), which face to the cytoplasm *in vivo*, share very low protein sequence identities (Fig. 6A). The estimated charge of the N-terminal domain of human TOM22 is strongly negative (−16) (Fig. 6B, upper), suggesting that the N-terminal domain captures the mitochondrial precursors via their positively charged TPs as shown in a recent study (Su et al., 2022). On the other hand, the estimated charge of the N-terminal domain of *C. merolae* TOM22 is not negative (+2.2), but we found that one putative α-helical domain has five acidic residues on one side (Fig. 6B, bottom). Thus, in *C. merolae*, mitochondrial precursor proteins would be verified via electrostatic residue-residue interaction between basic residues on the TP and the acidic residues on TOM22 of the α-helix as the single-step authentication (Fig. 6C). The TOM22-mediated authentication mechanism for mitochondrial protein import would be one reason why the TP with very few basic residues can fulfil the requirement for protein targeting to the mitochondrion in *C. merolae*.

**Figure 6.**
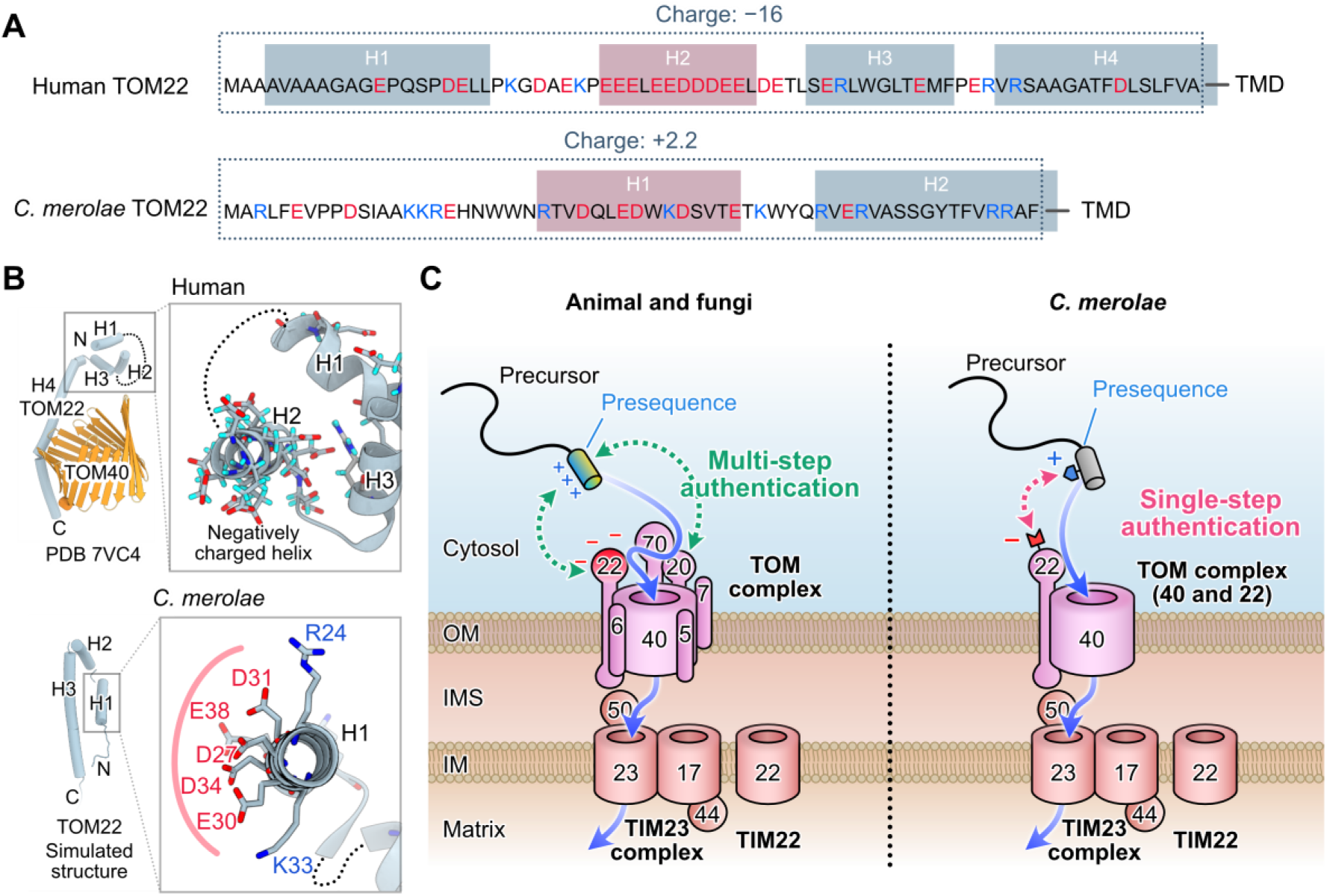
Schematic representation of mitochondrial protein import in *C. merolae*. (**A**) Comparison of the N-terminal domains of human TOM22 and *C. merolae* TOM22. Human TOM22 helix domains and *C. merolae* TOM22 helix domains are shown in boxes. Helical regions for *C. merolae* TOM22 are computationally predicted by NetSurfP2.0. Red boxes indicate negatively charged helices. (**B**) Protein architectures of TOM22 and TOM40. Human TOM22 and TOM40 are depicted by using the protein structural data (PDB:7VC4). Structure for *C. merolae* TOM22 was computationally simulated. Detail structures of boxed areas are shown in right side for each. (**C**) Schematic models for mitochondrial protein import in animal and fungi (left) and in *C. merolae* (right). In animal and fungi, a mitochondrial presequence, which is an amphiphilic helix is recogninzed by TOM20 and TOM22 as multi-step authentication process. In contrast, a preseuqnce α-helix containing a few numbers of basic residues would be recognized by the acidic side on the α-helix in TOM22 via electrostatic residue-residue interaction as single-step authentication in *C. merolae*. During the process, basic residues on preseuqnece work as key tooth and the acidic side on the TOM22 plays as the lock. After the authentication, a precursor protein would be imported into the TOM40 and drawn by the function of the TIM complex.

### Diversification of the TOM complex in eukaryotes during evolution

The requirement of arginine residues for protein targeting to mitochondria is also conserved in green plants (Heidorn-Czarna et al., 2022; Lee et al., 2019). However, the protein components of the TOM complex in green plants differ significantly not only from those in fungi/animals, but also from those in red algae (Carrie et al., 2010). In contrast to red algae, no orthologue of TOM22 has been identified in the green plant lineages. Furthermore, although *Arabidopsis thaliana* TOM20 has been shown to be functionally equivalent to animal/fungal TOM20, there is no sequence similarity between *A. thaliana* TOM20 and animal/fungal TOM20 (Perry et al., 2006). Thus, the TOM complex in the green plant lineage is composed of many types of specific proteins, but the authentication mechanism of mitochondrial precursor proteins in green plants could be similar in complexity to those in fungi and animals through convergent evolution. Considering the diversification of the components and the complexity of the structure of TOM complexes in modern eukaryotes, the simple authentication mechanism in the red alga *C. merolae* is very unique and would be a minimal or primitive system.

### Rewriting of the mitochondrial targeting information with an additional polypeptide for targeting to the chloroplast

Our results suggest a putative mechanism to classifying mitochondrial and chloroplast precursor proteins in *C. merolae*, which has the simplest cell structure as a photosynthetic eukaryote. Given that the mean length of the chloroplast TPs is longer than that of the mitochondrial TPs (Fig. 1E and 1F) and our experimental result of chloroplast RPSA (Fig. 5E), the presence of additional peptide sequences would determine whether the TP are for the mitochondrion or the chloroplast. The longer TP length for the chloroplast TP than that for the mitochondrial TP is also found in *Arabidopsis thaliana*. Since the chloroplast arose after the birth of the mitochondrion via endosymbiosis, the protein targeting system for the chloroplast would have to use longer, more complex TPs to distinguish it from the mitochondrial TP.

## Materials and Methods

### Cell culture

*C. merolae* M4 strain, which is isolated from *C. merolae* 10D (NIES-3377) and has a mutation in the *URA5.3* gene, was used in this study (Minoda et al., 2004). The *C. merolae* M4 cell were cultured in MA2 medium supplemented 0.5 mg/mL uracil and 0.8 mg/mL 5-fluoroorotic acid mono-hydrate in a tissue culture flask 25 (TPP Techno Plastid Products AG, Switzerland) with shaking at 120 rpm under continuous white light at 40°C.

### Putative presequence dataset

Protein sequences of well-studied mitochondrial and chloroplast proteins are chosen from 4803 *C. merolae* ORFs (Matsuzaki et al., 2004; Mori et al., 2016; Moriyama et al., 2014b; Moriyama et al., 2014a; Nozaki et al., 2007). By the protein-BLAST search, non-conserved region at the N-terminal of each protein sequence was identified by visual confirmation and the region is addressed as a putative presequence region in this study. Through the approach, we created the sequence dataset for 113 of mitochondrial presequences (Table S1) and 97 of chloroplast presequences (Table S2).

### Data analysis

Prediction of presequences for the mitochondrion and the chloroplast were performed by TargetP 2.0 program (Armenteros et al., 2019) (Data S1). Presequence net charge was calculated for a pH of 7 using the Protein Calculator v3.4 program (https://protcalc.sourceforge.net/). Hydrophobicity and hydrophobic moment of each putative α-helical polypeptide were calculated using the HELIQUEST program (Gautier et al., 2008). Amino acid frequencies were calculated using the *C. merolae* ORF data (http://czon.jp/). Sequence logos were made in the WebLogo program (Crooks et al., 2004). Helical wheel maps are made in the NetWheels program (http://lbqp.unb.br/NetWheels/). Protein local structures were predicted by the NetSurfP 2.0 program (Klausen et al., 2019) and tertial structures were simulated by the AlphaFold program (Jumper et al., 2021). Peptide sequence homologies between the synTP and N-terminal 1-24 amino acids of the *C. merolae* ORFs were calculated using BLOSUM30 matrix.

### Transformation and the fluorescent reporter assay

The *C. merolae* cell lines expressing mVenus fused with N-terminal targeting peptides (TP) were produced as follows. To generate the presequence fused mVenus expression vector, a presequence of interest was cloned into the vector by following methods. To produce the strains AAT (aspartate aminotransferase, CMC148C) 1-33, 3-R, 2-R, 1-R (R13), 0-R, 3-K, FMNL 1-24, FMNL 1-24 (E2A), FMNL 1-24 (E18A), FMNL 1-24 (E2A/E18A) and RPSA (chloroplast r-protein S1, CMM019C) 1-24, the nucleotide sequence of TPs were added to the 5’ end of *mVenus* by two-step PCR and cloned into the linearized pQE80L vector containing the *URA5.3* upstream sequence (from –2,300 to –898 bp), the 500 bp of *CpcC* promoter, the 200 bp of *TUBB* downstream sequence and the *URA5.3* selection marker. To produce the strains 1-R (R2), 1-R (R6), 1-R (R8), 1-R (R9), 1-R (R12), 1-R (R15), 1-R (R21), synthetic DNA fragments were clone into the linearized vector.

Using the resulting constructs as a template, DNA fragments for homologous recombination comprising ∼1400 bp of sequence upstream of URA5.3, the *CpcC* promoter, nucleotide sequence of TP, *mVenus*, the *TUBB* 3′ UTR and the *URA5.3* selection marker were amplified by PCR. The PCR amplicons were introduced into upstream of the chromosomal *ura5.3* of the uracil auxotrophic mutant strain M4 by polyethylene glycol (PEG)–mediated transformation as described previously (Fujiwara et al., 2015; Imamura et al., 2009; Ohnuma et al., 2008). Positive clones, which are non-uracil-auxotrophy, on the MA2 plate were confirmed by sanger sequencing and observed by fluorescence microscopy. See also Fig. S3.

All PCR amplification were performed with Platinum SuperFi II DNA polymerase (Thermo Fisher Scientific). Purification and assembly of DNA fragments were performed using a Wizard SV Gel and PCR Clean-up System (Promega) and a NEBuilder HiFi DNA assembly cloning kit (New England Biolabs). DNA sequences and resultant transformants are listed in Table S6, respectively.

### Fluorescence microscopy

Fluorescence observations were performed on an Olympus IX83 inverted microscope with a 1.45 NA, 100× oil immersion objective. Illumination was provided by a fluorescent light source (U-HGLGPS; Olympus), excitation filters [490-500HQ (Olympus) for mVenus, FF01-405/10-25 (Semrock) for chloroplasts], custom dichroic mirrors [Di03-R514-t1-25×36 (mVenus), Di03-R514-t1-25×36 (Semrock), T455lp (Chroma) (for chloroplasts)], emission filters [FF02-531/22-25 (Semrock) (for mVenus), FF02-617/73-25 (for chloroplasts) (Semrock)]. Images were acquired with a ORCA Fusion BT sCMOS camera (Hamamatsu Photonics) controlled by MetaMorph software (Molecular Devices). The effective pixel size was 65.2 nm x 65.2 nm.

### Phylogenetic analysis of *C. merolae* RPSA

The amino acid sequences of chloroplast r-protein S1 (CMM019C) and mitochondrial r-protein S1 (CMI304C) of *Cyanidioschyzon merolae* were retrieved from the *Cyanidioschyzon merolae* Genome Project v3 (Matsuzaki et al., 2004; Nozaki et al., 2007) and used as a query for BLASTP (http://blast.ncbi.nlm.nih.gov/Blast.cgi). The BLASTP was carried out against the non-redundant protein sequences (nr) of National Center for Biotechnology Information database (NCBI, http://www.ncbi. nlm.nih.gov/) all organisms. The top 50 sequences with the highest E-value sequences of each query were retrieved. We also added amino acid sequences annotated with chloroplast r-protein S1 of six Viridiplantae species from NCBI. Multiple sequence alignments were generated using MAFFT v7.511 online service in auto strategy (Katoh et al., 2018; Kuraku et al., 2013). Non-homologous regions were detected and cleaned using HMMcleaner (Di Franco et al., 2019). Finally, the multiple sequence alignment was trimmed using trimAl by automated trimming heuristic using “-automated1” option (Capella-Gutiérrez et al., 2009). Redundant OTUs that became identical after the trimming procedures were manually excluded. Finally, 83 OTUs were used for analysis. Bayesian inference for the alignments was carried out using MrBayes v3.2.7a MPI version with the best-fitted model selected by ModelTest-NG v0.1.7 (Altekar et al., 2004; Darriba et al., 2020; Flouri et al., 2015). Convergences of Markov chain Monte Carlo iterations were evaluated based on the average standard deviation of split frequencies for every 1,000,000 generations, discarding the first 25% as burn-in, and the iterations were automatically stopped when the average standard deviations were below 0.01, indicating convergence. In addition, the maximum likelihood method was subjected to the alignment with bootstrap values based on 1,000 replications by RAxML-NG v1.2.0 with the same model as Bayesian inference (Kozlov et al., 2019). The alignments used for the phylogenetic analysis are uploaded to TreeBASE (Vos et al., 2012).

## Acknowledgements

We thank our lab colleagues for their support and advice during this project.

## Author Contributions

Conceptualization: R.H., Y.M., Y.Y.; Methodology: R. H., Y. M., K.T., H. N., Y.Y.; Investigation: R.H., Y.M. K.T., Y.Y.; Writing – original draft: R.H, Y.M., Y.Y.; Writing – review & editing: R.H., Y.M., K.T., H.N., T.H., Y.Y.; Visualization: R.H., Y.Y.; Supervision: Y.Y.; Project administration: Y.Y.; Funding acquisition: Y.Y.

## Data availability

The alignments used for the phylogenetic analysis are available in TreeBASE under the following accession codes: S31160. The data supporting the findings of this work are available in the main figures and supplementary information.

---

**Figure S1.**
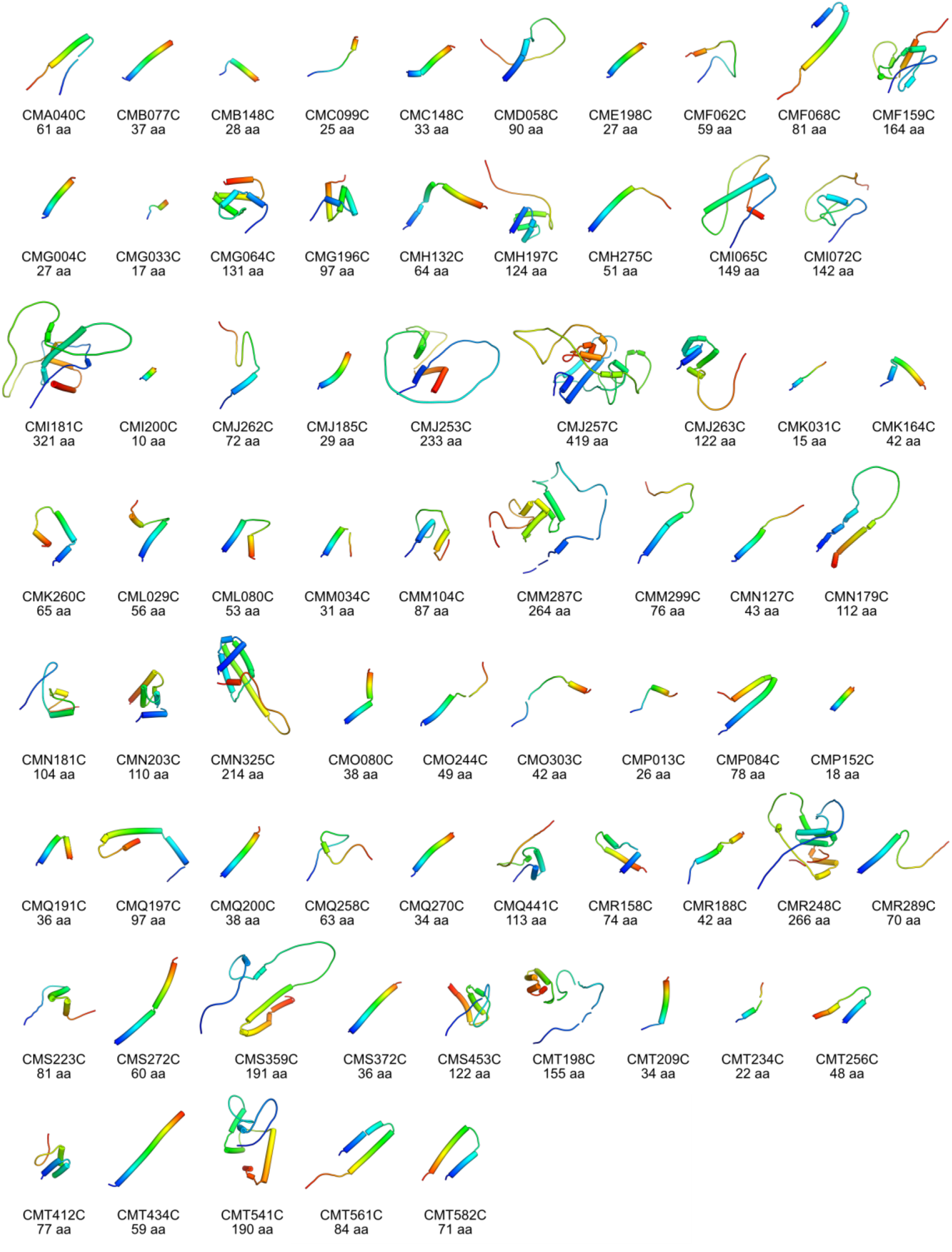
Simulated structures of mitochondrial presequences containing α-helical regions. In the 113 mitochondrial presequences, 71 presequences contain α-helical structures as a secondary structure. Tertial protein models of mitochondrial presequences are rainbow-colored from the N (blue) to the C terminus (red).

**Figure S2.**
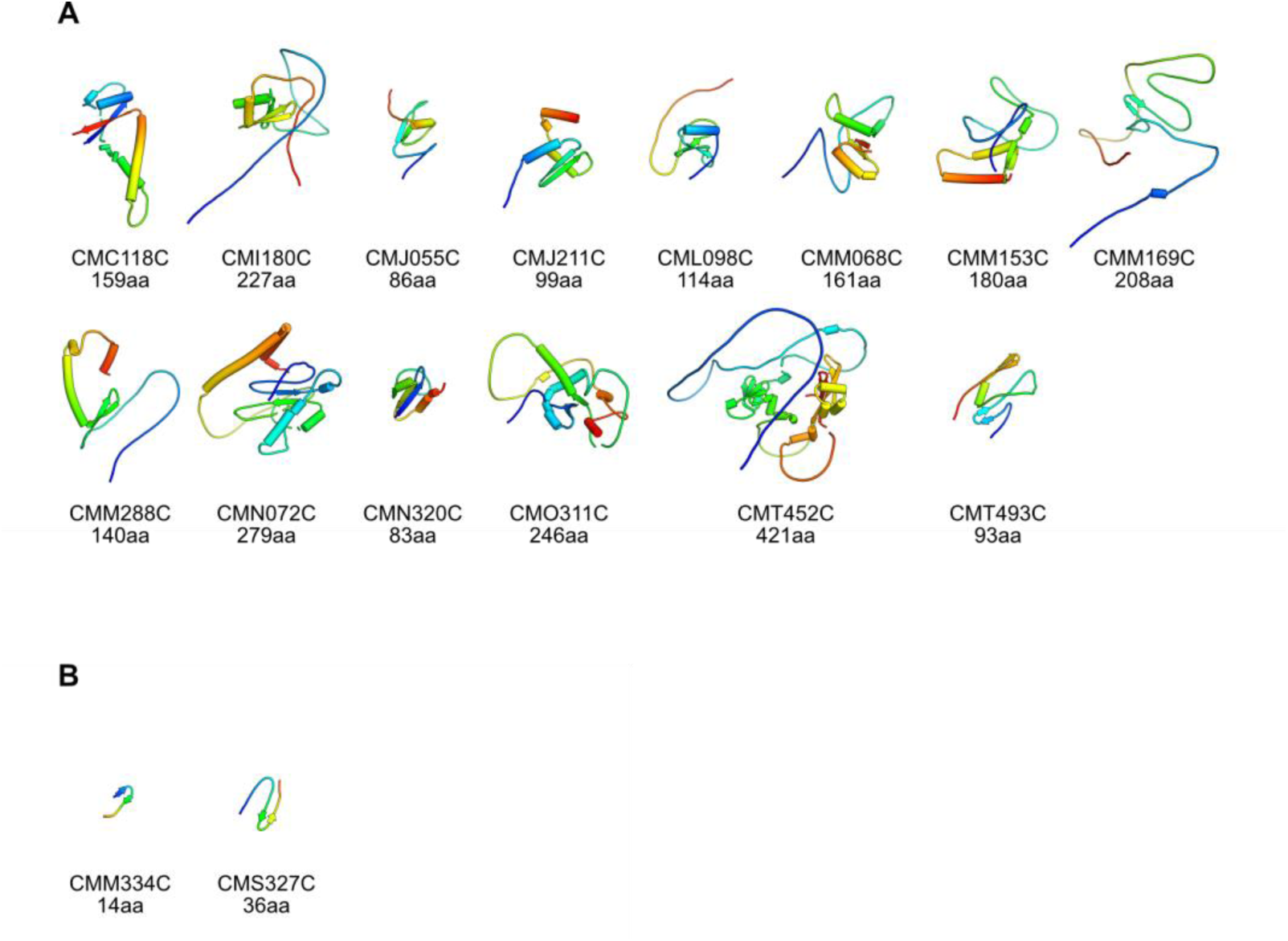
Simulated structures of mitochondrial presequences containing α-helical and β-sheet regions. (**A** and **B**) Presequences containing both α-helix and β-sheet regions (**A**) and only β-sheet regions (**B**). In the 113 mitochondrial presequences, 14 presequences contain both α-helical and β-sheet structures and 2 presequences contain only β-sheet structures. Tertial protein models of mitochondrial presequences are rainbow-colored from the N (blue) to the C terminus (red).

**Figure S3.**
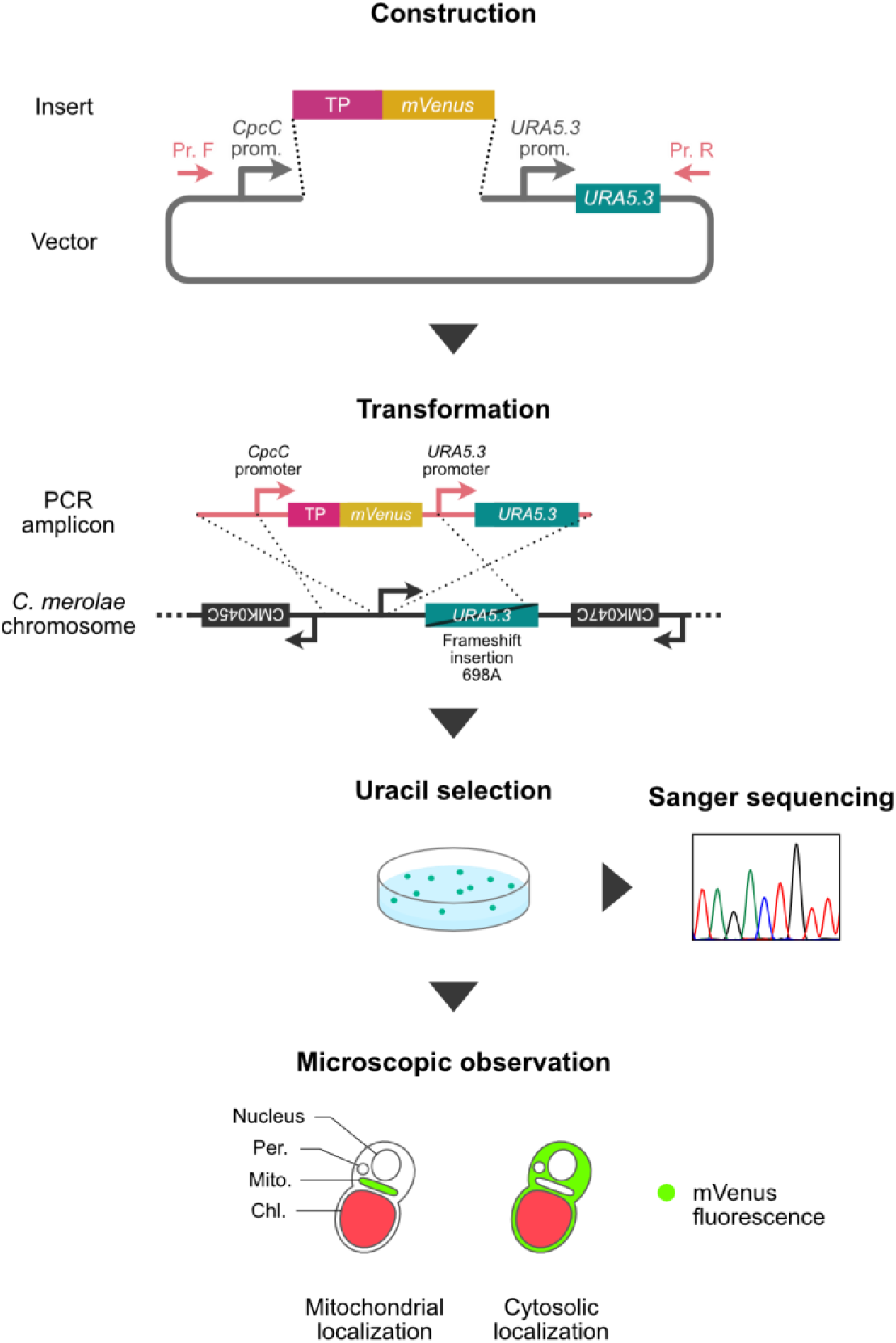
Schematic representation of the fluorescent reporter assay for evaluation of the protein targeting to the mitochondrion. Further details on the construction of the plasmids and fluorescence microscopy are described in the Materials and Methods section.

**Figure S4.**
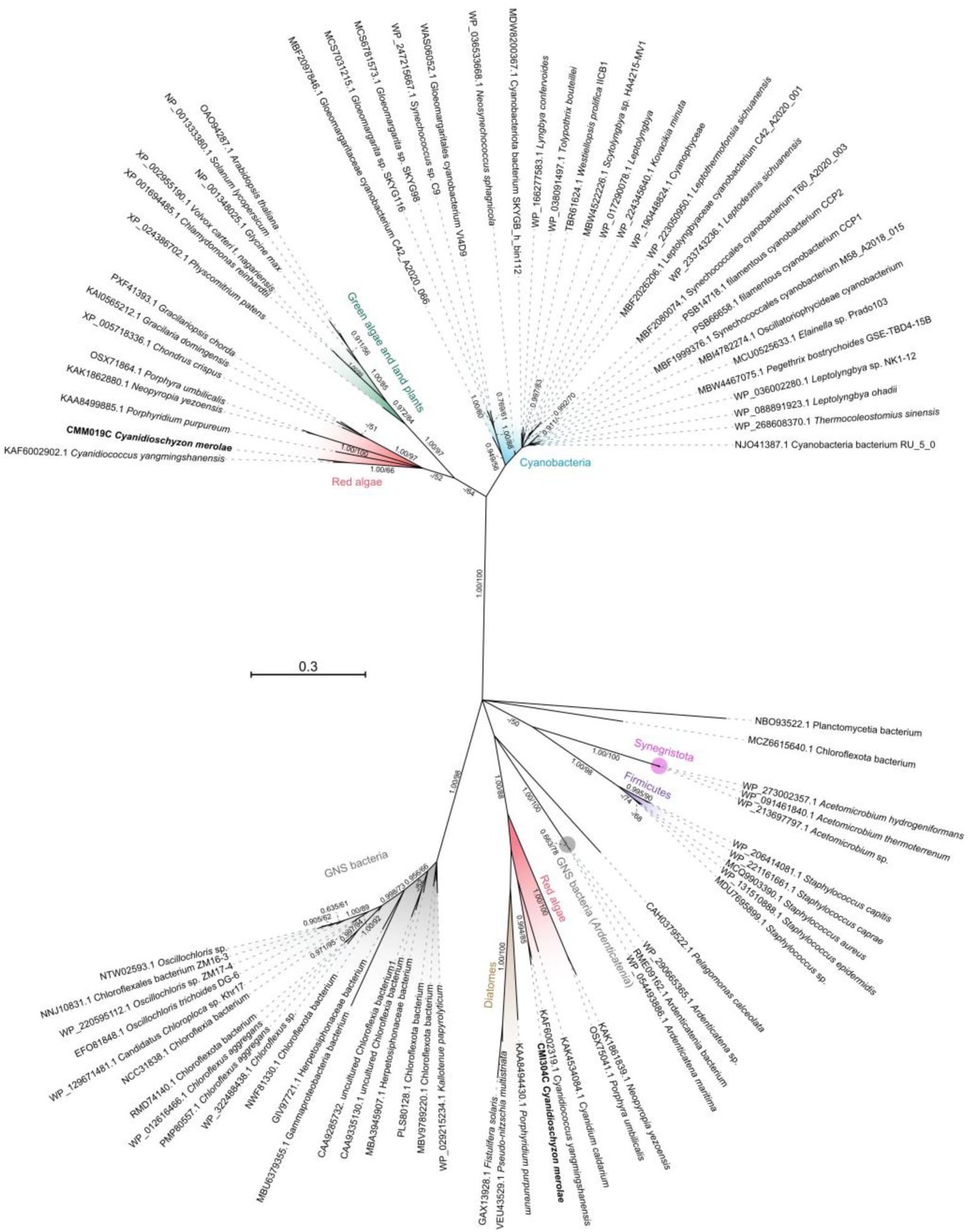
The unabbreviated Bayesian tree of chloroplast, mitochondrial and bacterial RPSA proteins. Phylogenetic analysis of *C. merolae* chloroplast and mitochondrial RPSAs (CMM019C and CMI304C), inferred based on 226 amino acid sequences by the Bayesian inference (BI) using the LG + G4 model. The number before the species name of each OTU indicates the NCBI accession number for the sequence. Branch lengths are proportional to the evolutionary distances indicated by the scale bar. Numbers at left and right above branches indicate posterior probabilities of BI (≥0.90) and bootstrap values of the maximum likelihood method (≥50%), respectively.

**Table S1.**
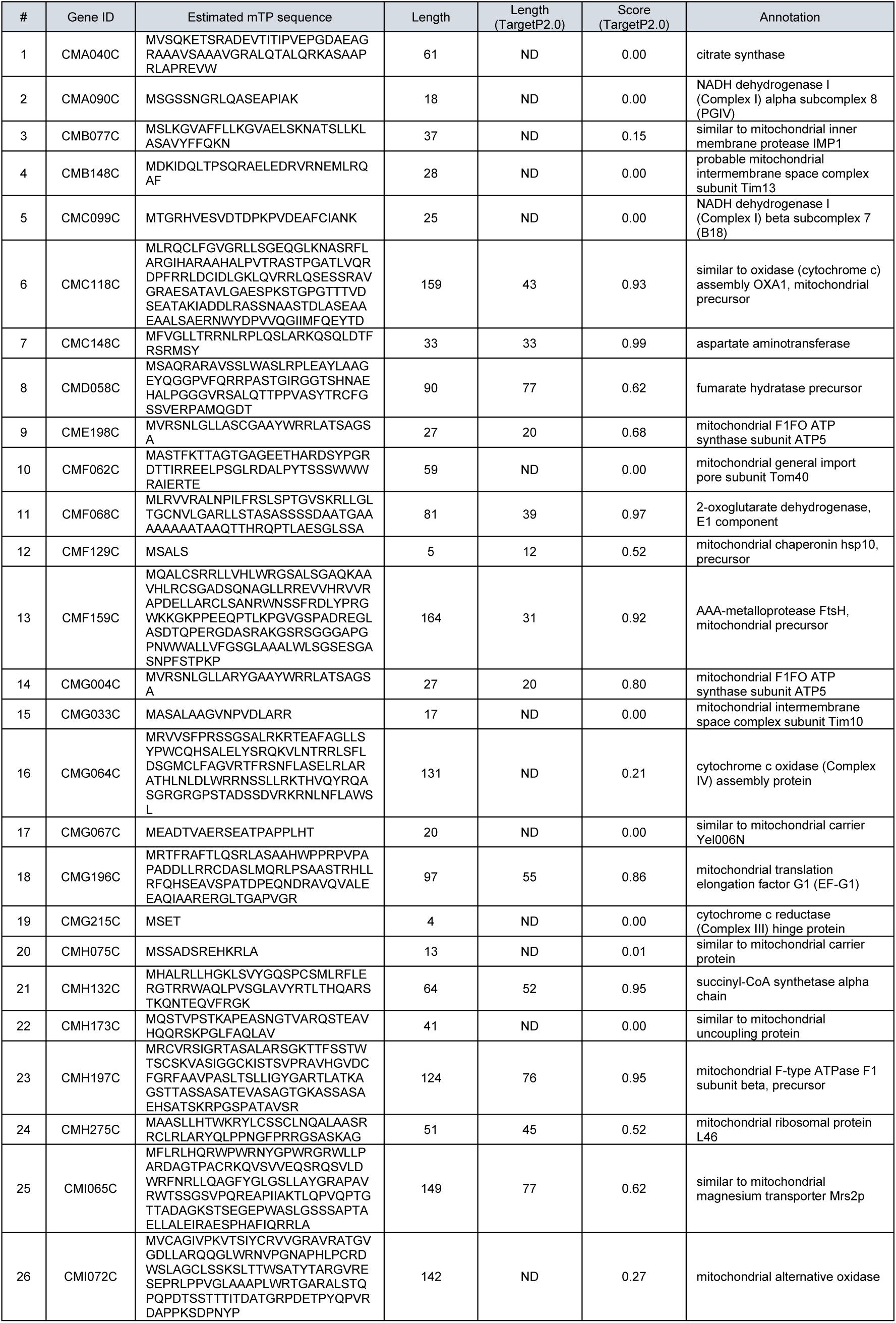
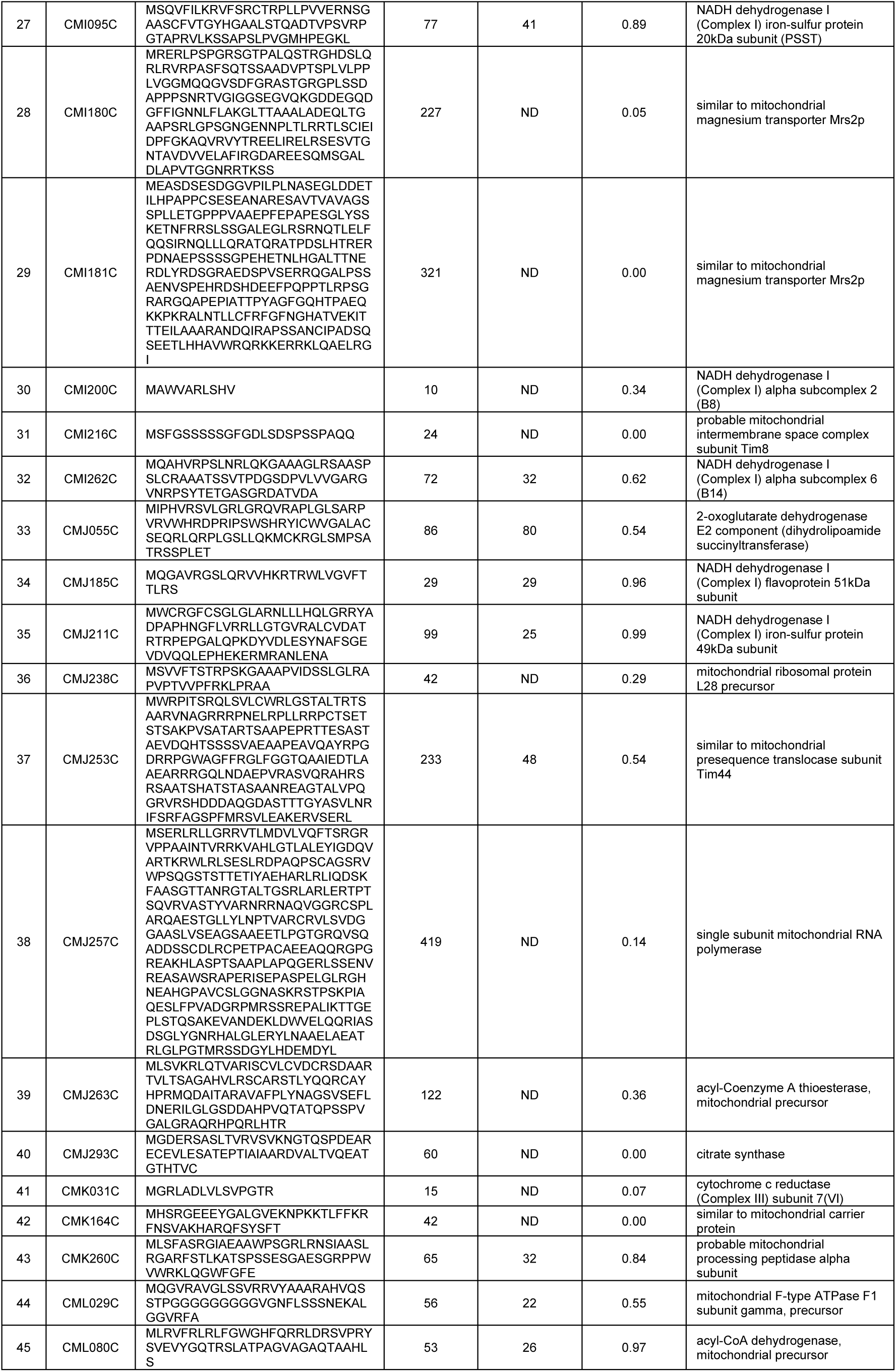
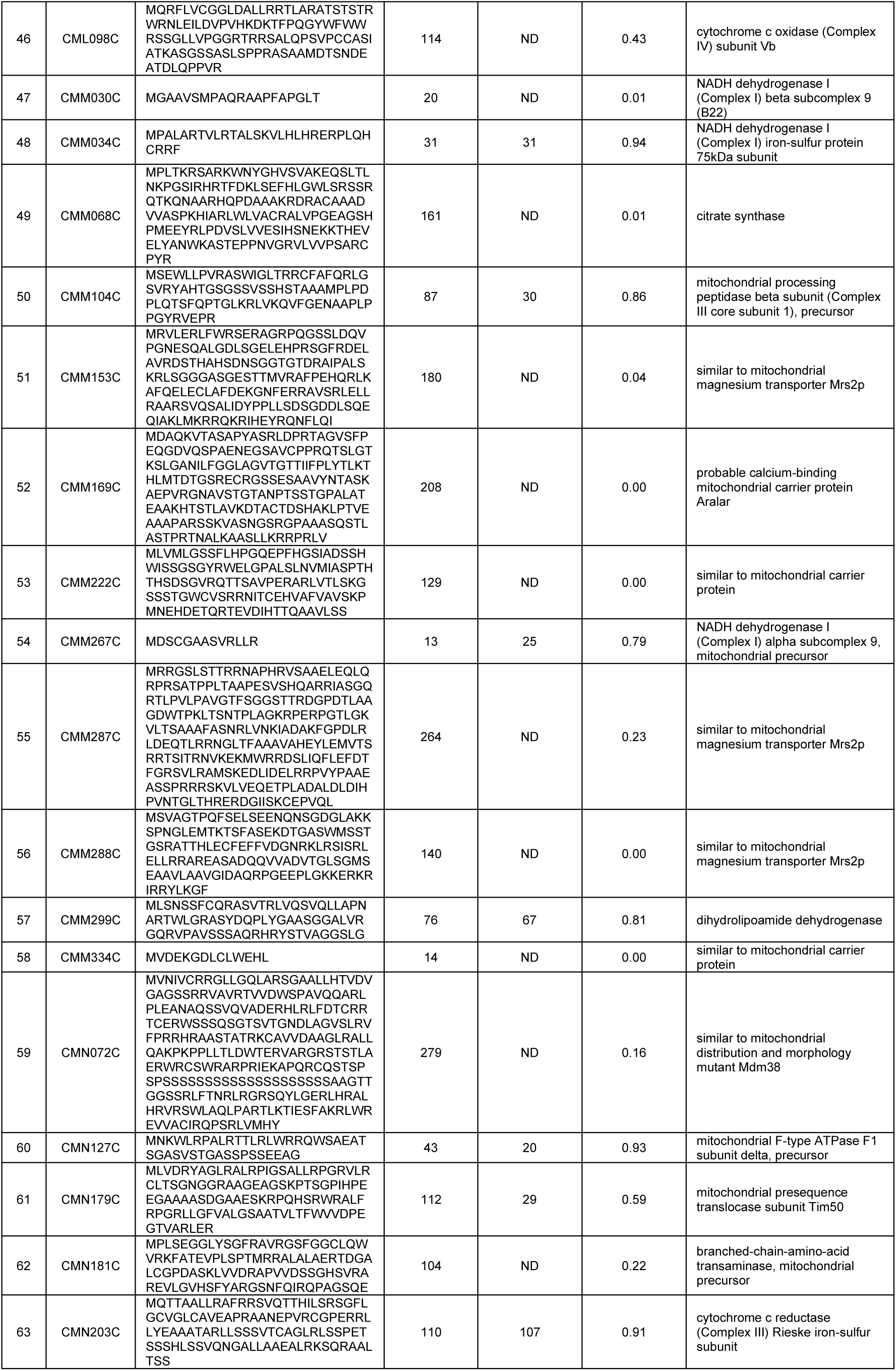
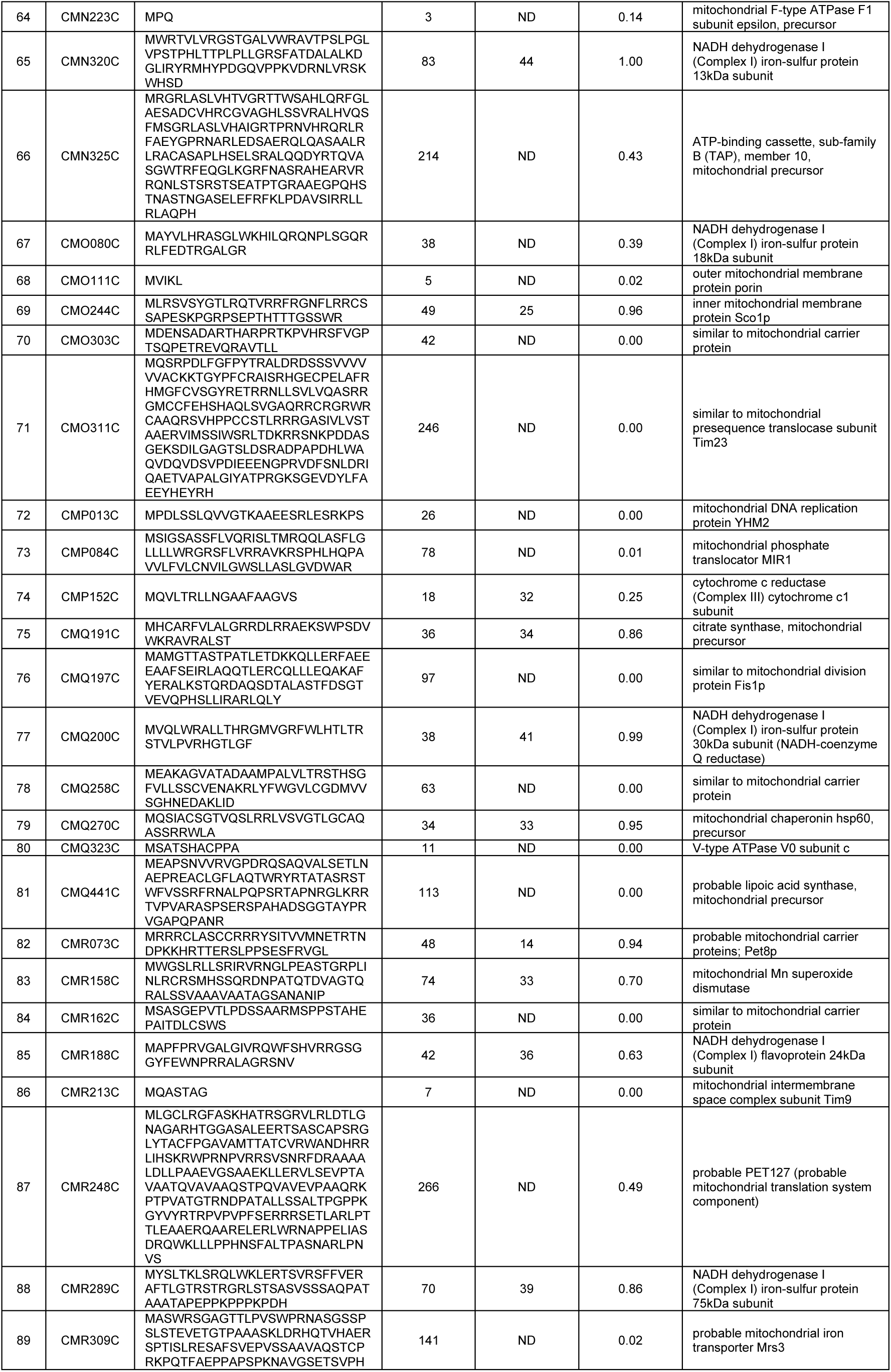
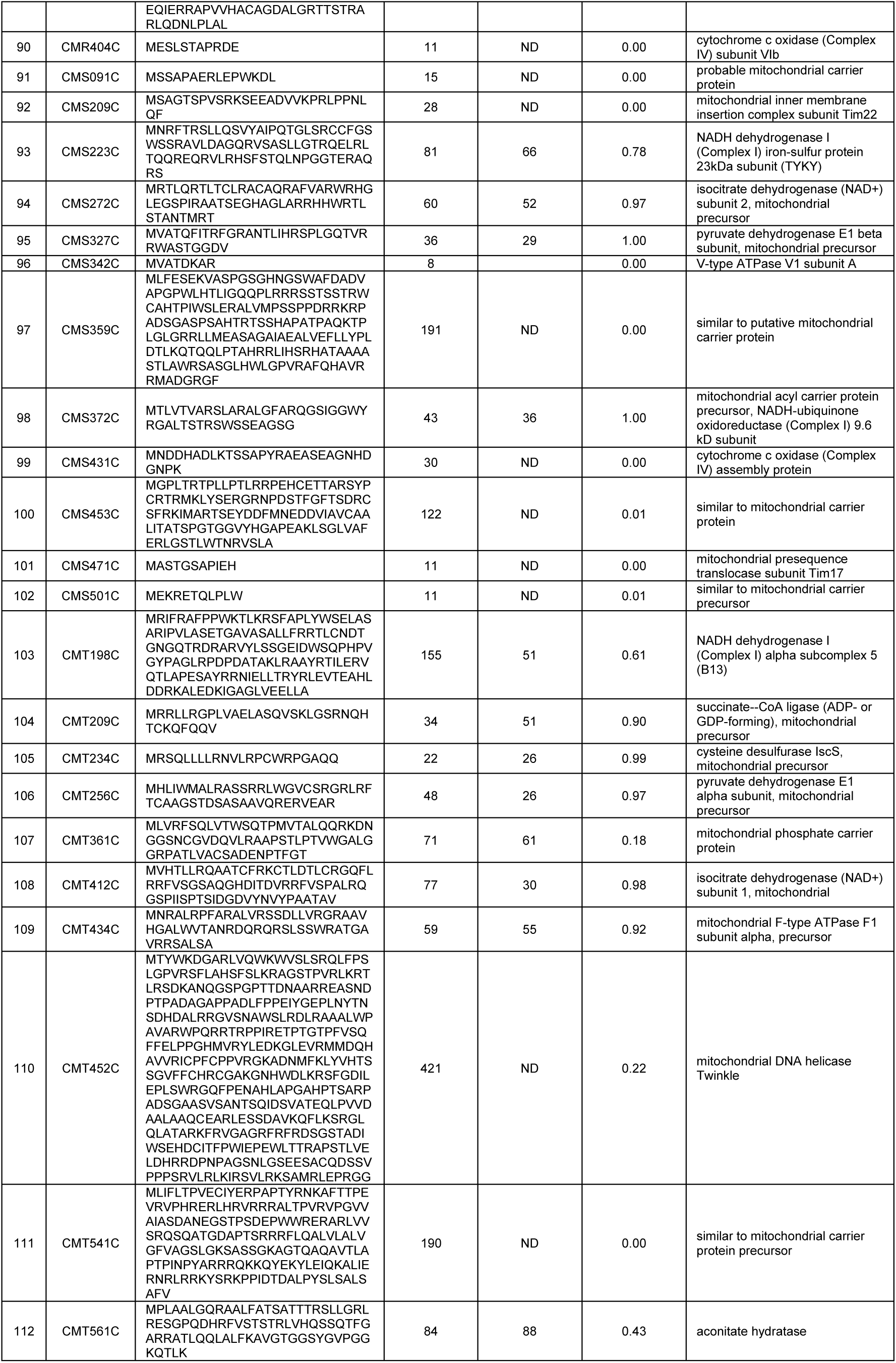
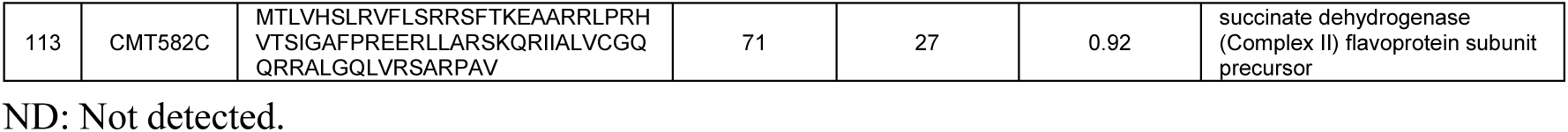
List of 113 putative mitochondrial TP sequences.

**Table S2.**
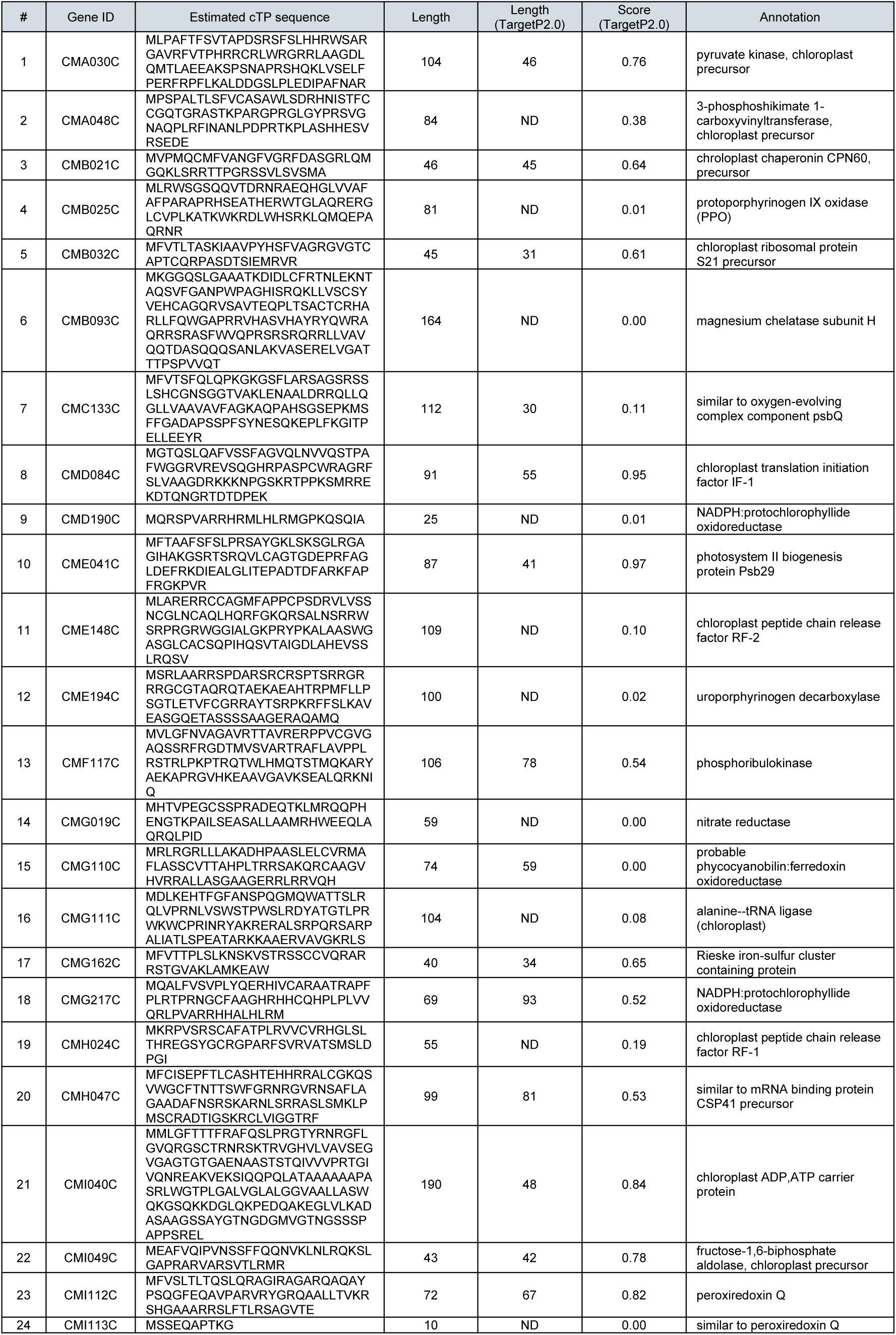
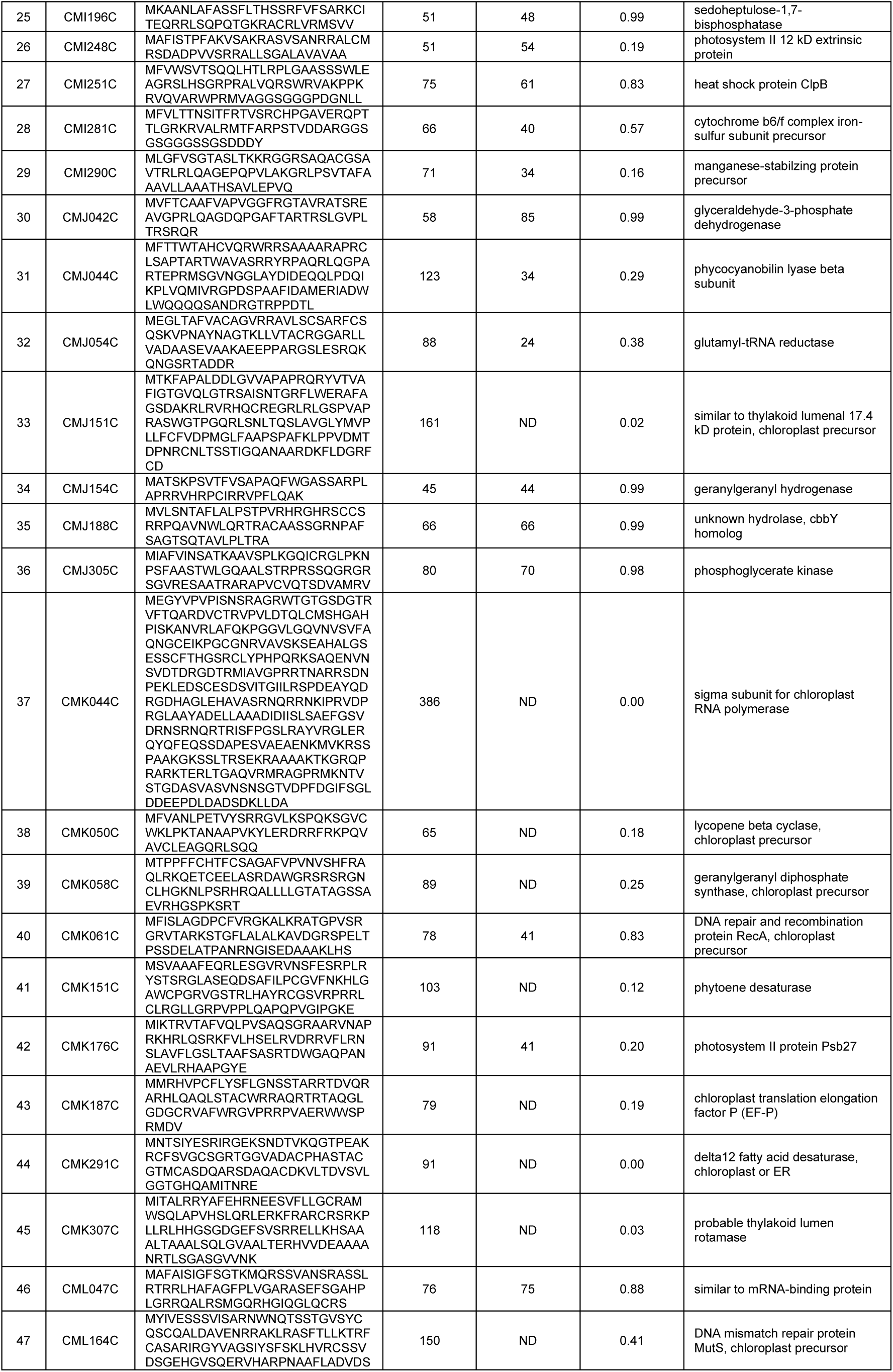
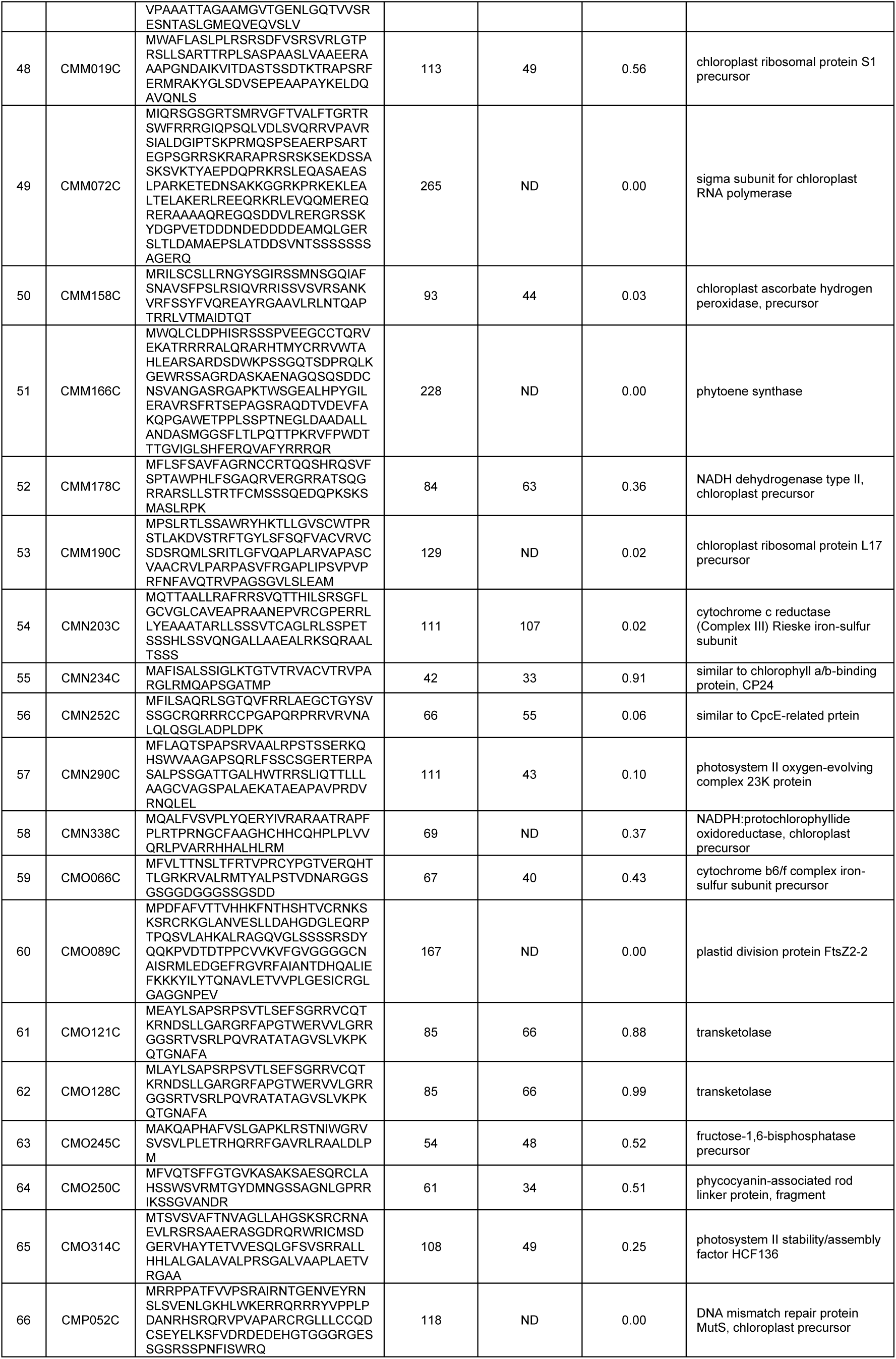
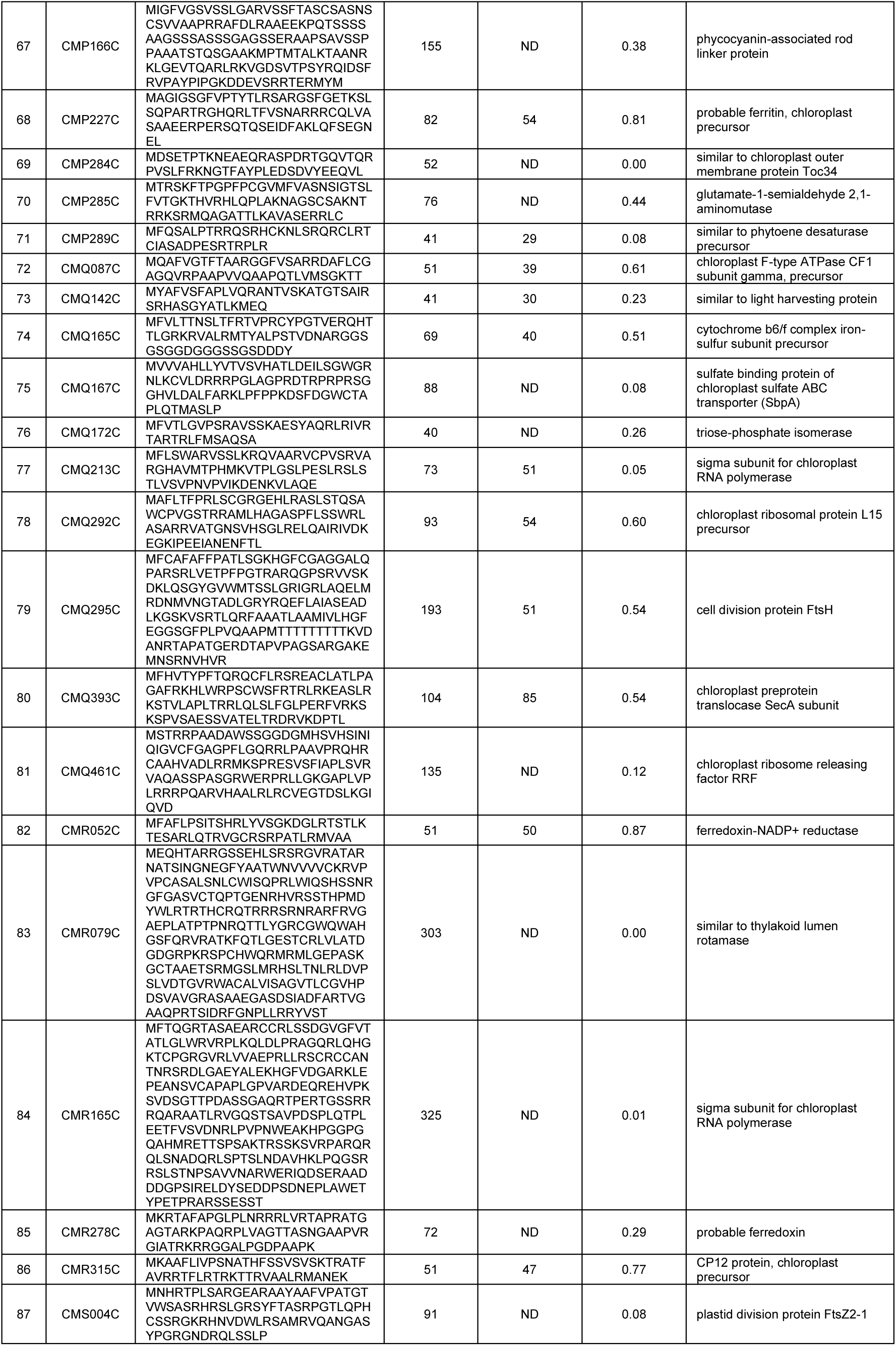
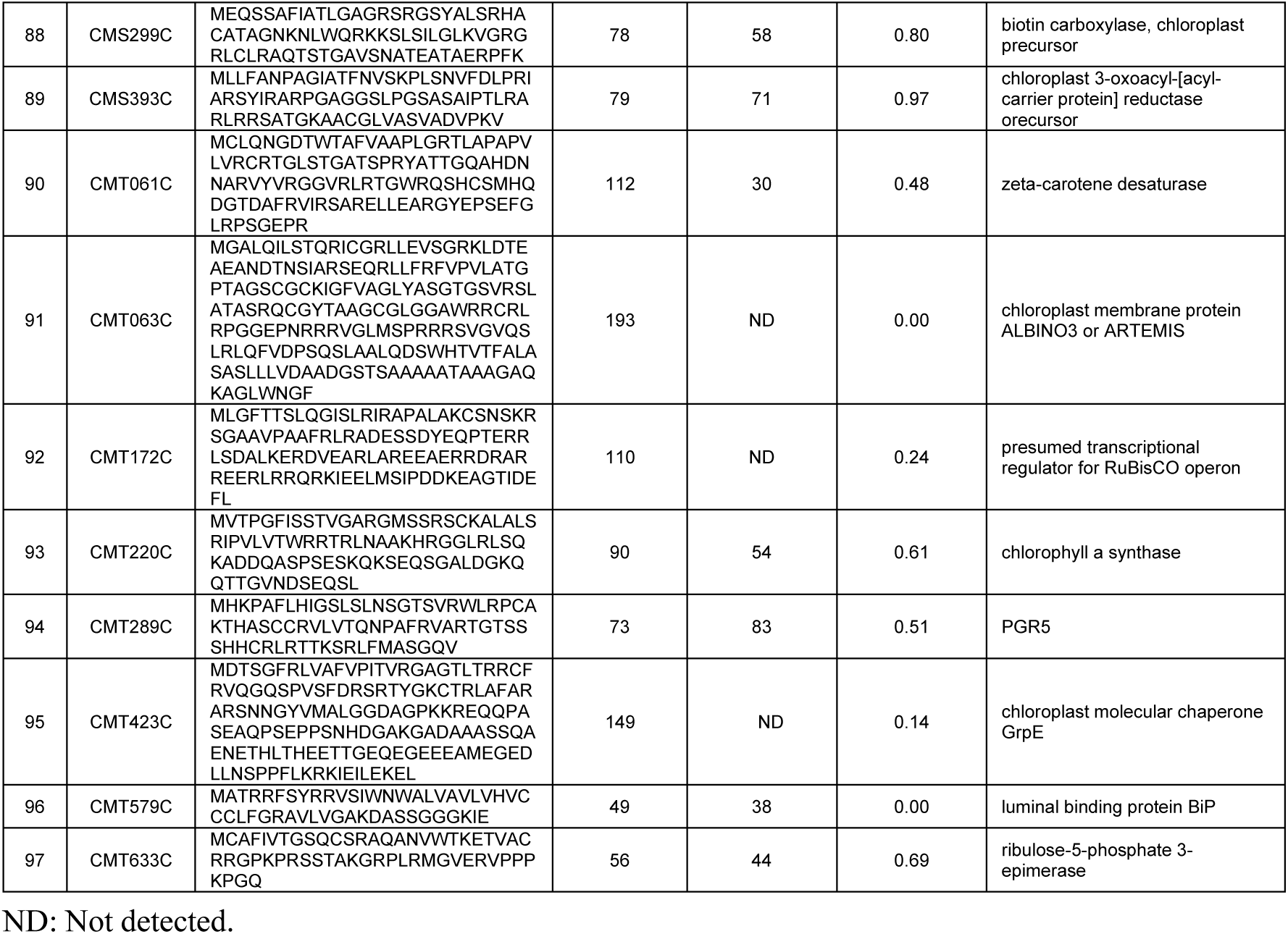
List of 97 putative chloroplast TP sequences.

**Table S3.**
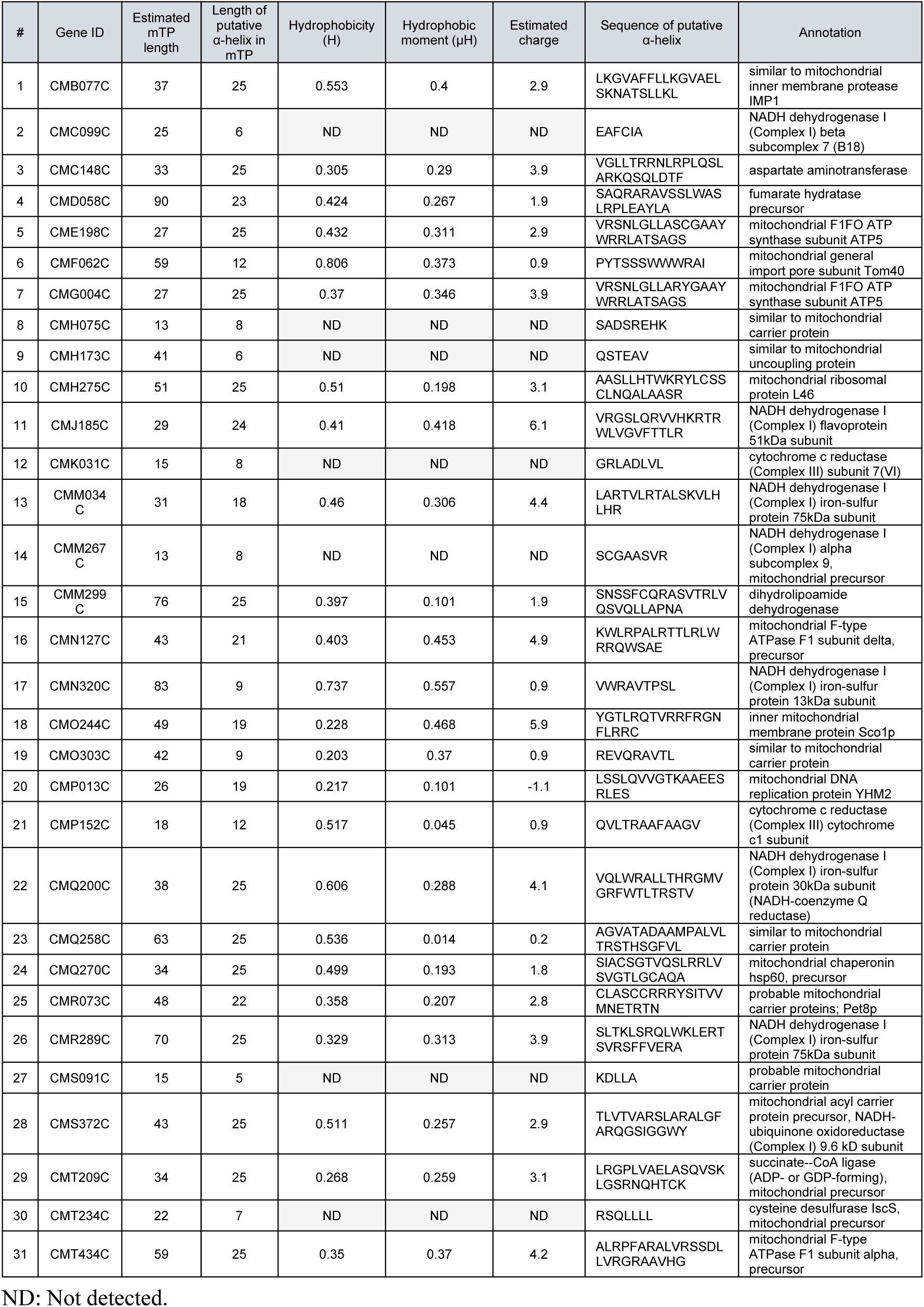
List of putative α-helical sequences in mitochondrial TPs.

**Table S4.** Amino acid sequence and relative scores of each synthetic TP.

| Name | Position |  |  |  |  |  |  |  |  |  |  |  |  |  |  |  |  |  |  |  |  |  |  |  | TargetP2.0 prediction results |  | Experimental result (this study) |
| --- | --- | --- | --- | --- | --- | --- | --- | --- | --- | --- | --- | --- | --- | --- | --- | --- | --- | --- | --- | --- | --- | --- | --- | --- | --- | --- | --- |
|  | 1 | 2 | 3 | 4 | 5 | 6 | 7 | 8 | 9 | 10 | 11 | 12 | 13 | 14 | 15 | 16 | 17 | 18 | 19 | 20 | 21 | 22 | 23 | 24 | mTP Score | SP Score |  |
| 3-R | M | S | A | L | L | L | A | L | L | L | S | L | R | A | S | L | L | R | R | T | A | T | A | A | 0.726 | 0.246 | Mitochondrion |
| 2-R | M | S | A | L | L | L | A | L | L | L | S | L | R | A | S | L | L | V | R | T | A | T | A | A | 0.342 | 0.649 | Mitochondrion |
| 1-R (R2) | M | R | A | L | L | L | A | L | L | L | S | L | A | A | S | L | L | V | E | T | A | T | A | A | 0.000 | 1.000 | cytosol |
| 1-R (R3) | M | S | R | L | L | L | A | L | L | L | S | L | A | A | S | L | L | V | E | T | A | T | A | A | 0.000 | 1.000 | NT |
| 1-R (R4) | M | S | A | R | L | L | A | L | L | L | S | L | A | A | S | L | L | V | E | T | A | T | A | A | 0.000 | 1.000 | NT |
| 1-R (R5) | M | S | A | L | R | L | A | L | L | L | S | L | A | A | S | L | L | V | E | T | A | T | A | A | 0.000 | 1.000 | NT |
| 1-R (R6) | M | S | A | L | L | R | A | L | L | L | S | L | A | A | S | L | L | V | E | T | A | T | A | A | 0.000 | 1.000 | Mitochondrion |
| 1-R (R7) | M | S | A | L | L | L | R | L | L | L | S | L | A | A | S | L | L | V | E | T | A | T | A | A | 0.001 | 0.999 | NT |
| 1-R (R8) | M | S | A | L | L | L | A | R | L | L | S | L | A | A | S | L | L | V | E | T | A | T | A | A | 0.009 | 0.990 | Mitochondrion |
| 1-R (R9) | M | S | A | L | L | L | A | L | R | L | S | L | A | A | S | L | L | V | E | T | A | T | A | A | 0.045 | 0.947 | Mitochondrion |
| 1-R (R10) | M | S | A | L | L | L | A | L | L | R | S | L | A | A | S | L | L | V | E | T | A | T | A | A | 0.151 | 0.825 | NT |
| 1-R (R11) | M | S | A | L | L | L | A | L | L | L | R | L | A | A | S | L | L | V | E | T | A | T | A | A | 0.004 | 0.993 | NT |
| 1-R (R12) | M | S | A | L | L | L | A | L | L | L | S | R | A | A | S | L | L | V | E | T | A | T | A | A | 0.090 | 0.869 | Mitochondrion |
| 1-R (R13) | M | S | A | L | L | L | A | L | L | L | S | L | R | A | S | L | L | V | E | T | A | T | A | A | 0.019 | 0.971 | Mitochondrion |
| 1-R (R14) | M | S | A | L | L | L | A | L | L | L | S | L | A | R | S | L | L | V | E | T | A | T | A | A | 0.010 | 0.985 | NT |
| 1-R (R15) | M | S | A | L | L | L | A | L | L | L | S | L | A | A | R | L | L | V | E | T | A | T | A | A | 0.003 | 0.996 | Mitochondrion |
| 1-R (R16) | M | S | A | L | L | L | A | L | L | L | S | L | A | A | S | R | L | V | E | T | A | T | A | A | 0.003 | 0.996 | NT |
| 1-R (R17) | M | S | A | L | L | L | A | L | L | L | S | L | A | A | S | L | R | V | E | T | A | T | A | A | 0.001 | 0.999 | NT |
| 1-R (R18) | M | S | A | L | L | L | A | L | L | L | S | L | A | A | S | L | L | R | E | T | A | T | A | A | 0.001 | 0.998 | NT |
| 1-R (R19) | M | S | A | L | L | L | A | L | L | L | S | L | A | A | S | L | L | V | R | T | A | T | A | A | 0.000 | 1.000 | NT |
| 1-R (R20) | M | S | A | L | L | L | A | L | L | L | S | L | A | A | S | L | L | V | E | R | A | T | A | A | 0.000 | 1.000 | NT |
| 1-R (R21) | M | S | A | L | L | L | A | L | L | L | S | L | A | A | S | L | L | V | E | T | R | T | A | A | 0.000 | 1.000 | Mitochondrion |
| 1-R (R22) | M | S | A | L | L | L | A | L | L | L | S | L | A | A | S | L | L | V | E | T | A | R | A | A | 0.000 | 1.000 | NT |
| 1-R (R23) | M | S | A | L | L | L | A | L | L | L | S | L | A | A | S | L | L | V | E | T | A | T | R | A | 0.000 | 1.000 | NT |
| 1-R (R24) | M | S | A | L | L | L | A | L | L | L | S | L | A | A | S | L | L | V | E | T | A | T | A | R | 0.000 | 1.000 | NT |
| 0-R | M | S | A | L | L | L | A | L | L | L | S | L | A | A | S | L | L | V | E | T | A | T | A | A | 0.000 | 1.000 | cytosol |
| 3-K | M | S | A | L | L | L | A | L | L | L | S | L | K | A | S | L | L | K | K | T | A | T | A | A | 0.099 | 0.848 | Mitochondrion |

NT: Not tested.

**Table S5.**
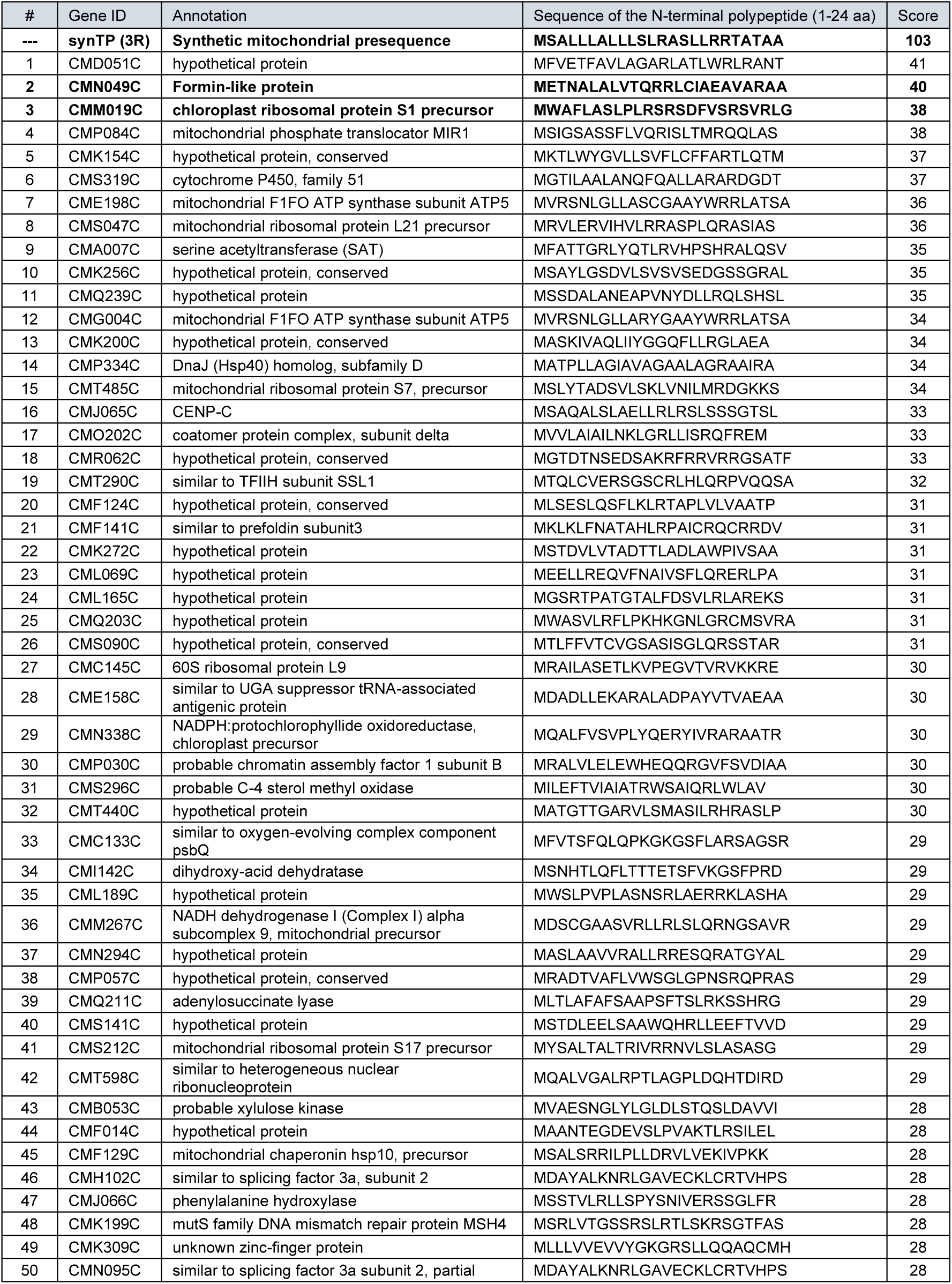
Top 50 of the peptide sequences that are similar to the synthetic TP evaluated by BLOSUM30.

**Table S6.**
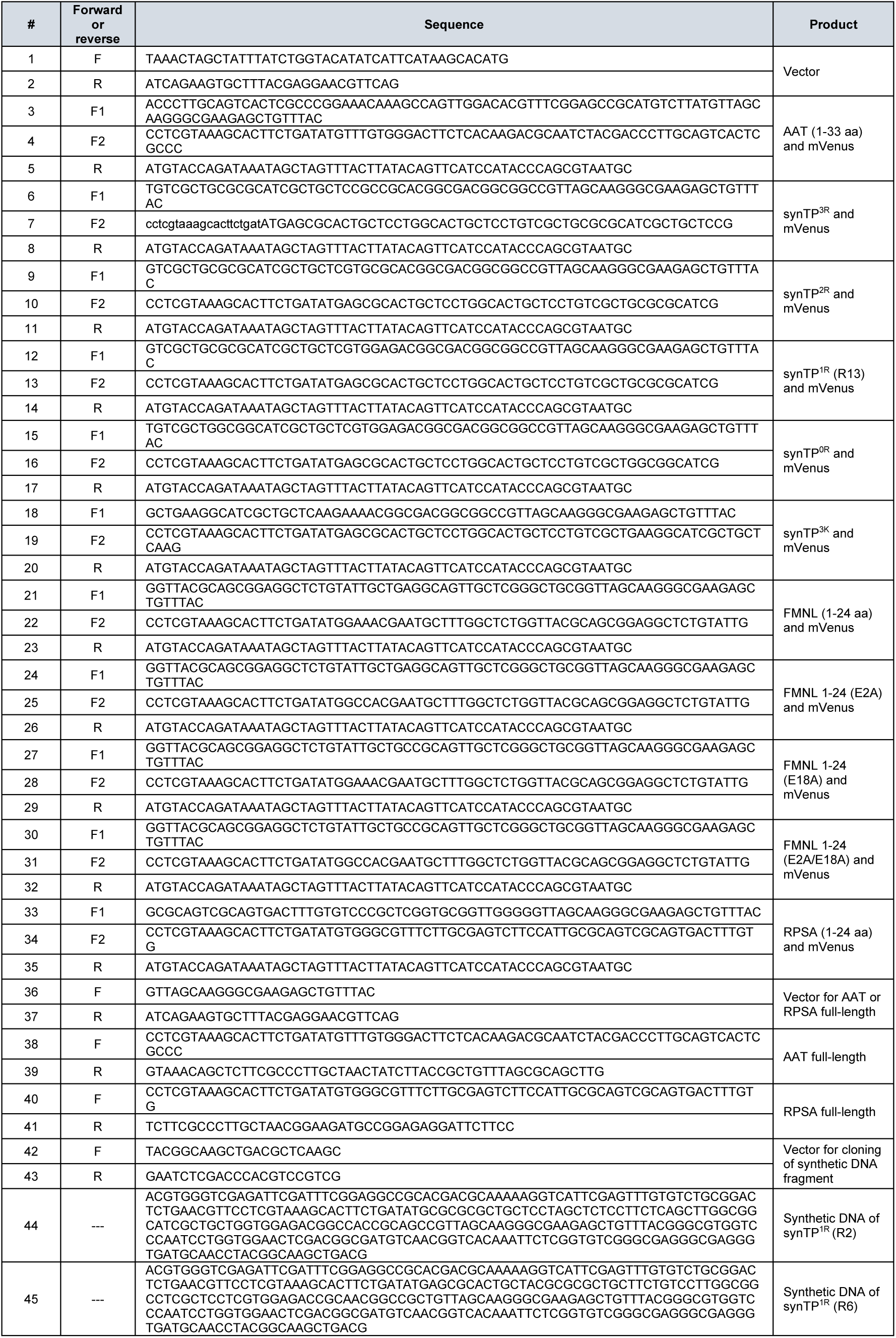
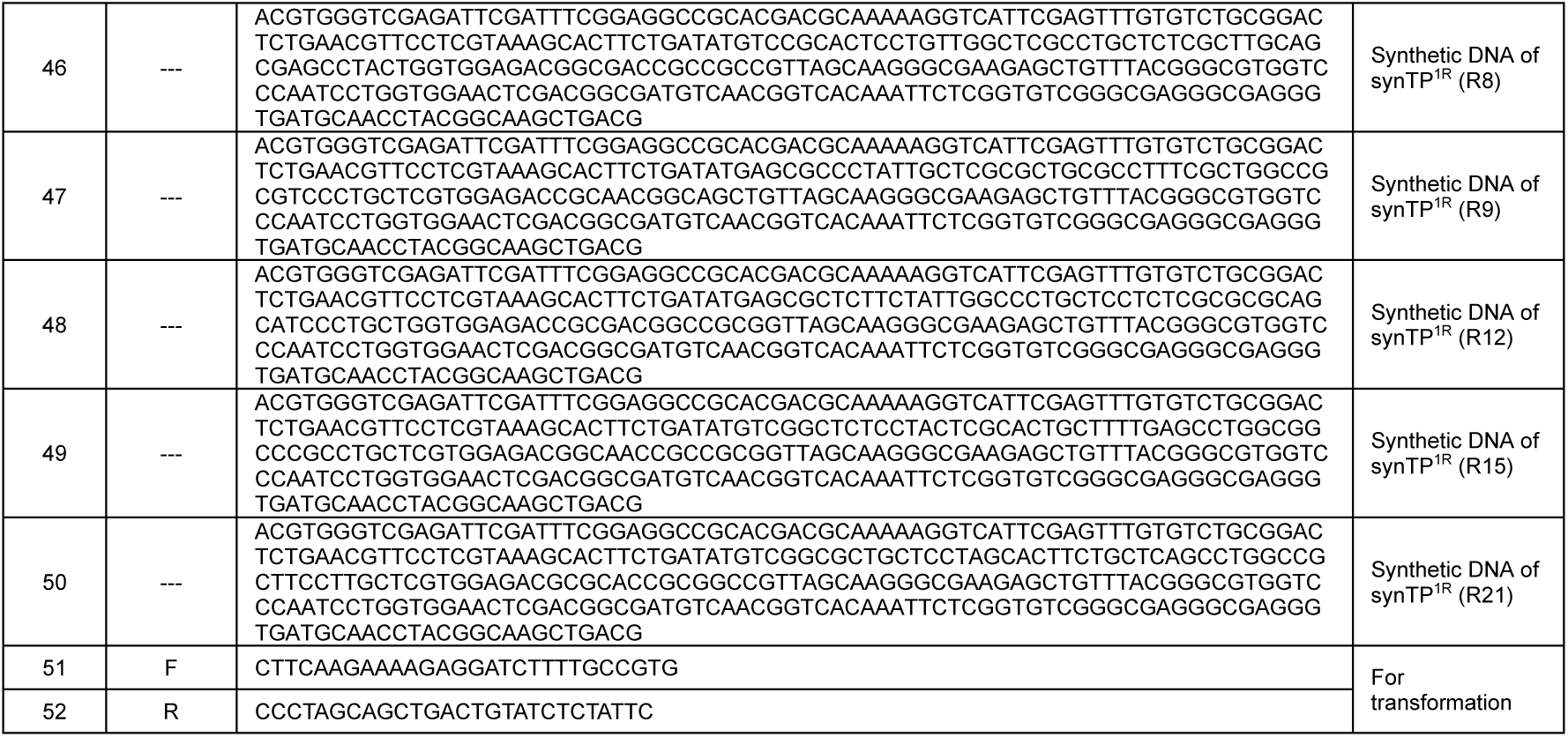
Primers and synthetic DNA fragments used in this study.

